# 5-HT1A Antagonist WAY-100635 Prevents Chronic Neurodegeneration and Immune Activation After TBI

**DOI:** 10.64898/2026.10.11.758371

**Authors:** Sayanta Dutta, Lilesh Kumar Pradhan, Jie Chen, Nian Wang, Xiang Gao, Arupratan Das

**Affiliations:** Department of Ophthalmology, Eugene and Marilyn Glick Eye Institute, Indiana University School of Medicine, Indianapolis, IN, USA; Department of Neurological Surgery, Indiana University School of Medicine, Indianapolis, IN, USA; Spinal Cord and Brain Injury Research Group, Stark Neurosciences Research Institute, Indianapolis, IN, USA; Advanced Imaging Research Center, The University of Texas Southwestern Medical Center, Dallas, Texas, USA; Department of Biomedical Engineering, University of Texas Southwestern Medical Center, Dallas, Texas, USA; Peter O’Donnell Jr. Brain Institute, University of Texas Southwestern Medical Center, Dallas, Texas, USA; Department of Medical and Molecular Genetics, Indiana University School of Medicine, Indianapolis, IN, USA; Stark Neurosciences Research Institute, Indiana University School of Medicine, Indianapolis, IN, USA

**Keywords:** Traumatic brain injury, Retinal ganglion cells, Neuroprotection, 5-HT1A receptor, Neuroinflammation

## Abstract

Traumatic brain injury (TBI) initiates secondary neurodegeneration that compromises interconnected neural circuits and contributes to persistent visual and neurobehavioral impairment. Pharmacological strategies that preserve vulnerable neurons and synapses while limiting chronic neuroinflammation remain an unmet therapeutic need. We previously identified the serotonin 1A receptor (5-HT1A) antagonist WAY-100635 through screening human stem cell-derived retinal ganglion cells (RGCs) and demonstrated that it promotes neuroprotection through cyclic adenosine monophosphate-dependent mitochondrial biogenesis. Here, we investigated whether post-injury treatment extends this neuroprotective efficacy to cerebral and visual circuitry following TBI.

Male C57BL/6J mice aged 2-3 months underwent unilateral controlled cortical impact (CCI) involving the left visual cortex. Mice were randomly assigned to sham, untreated injury or treated injury groups, with 3-10 sham controls and 5-8 mice per injury group across experiments. WAY-100635 was administered intraperitoneally at 5 mg/kg daily, beginning 1 hour after injury and continuing for 6 weeks. Therapeutic efficacy was assessed using motor and psychological-health-relevant risk-taking/behavioral disinhibition by elevated-plus-maze assessment, visual electrophysiology, optokinetic responses, diffusion magnetic resonance imaging, and retinal and cortical histopathology.

Unilateral cortical injury produced bilateral RGC dysfunction and loss, with greater degeneration in the eye contralateral to the injured hemisphere, consistent with predominantly crossed murine visual projections. WAY-100635 significantly preserved RGC survival and function, improved visual acuity and contrast sensitivity, and preserved visually evoked cortical responses to contralateral eye stimulation. Diffusion MRI revealed bilateral visual pathway abnormalities, with treatment selectively normalizing fractional anisotropy and radial kurtosis in the optic tract ipsilateral to cortical injury. In perilesional cortex, treatment preserved neuronal cell bodies and synapses detected by synaptophysin immunoreactivity while markedly reducing microglial/macrophage activation and reactive astrogliosis. These effects were accompanied by improved early motor performance and attenuation of injury-associated increases in open-arm exploration in the elevated plus maze. Cortical cavity volume was not significantly reduced, supporting preferential protection of vulnerable circuitry surrounding and connected to the primary lesion.

These findings identify post-injury WAY-100635 treatment as a novel therapeutic intervention that combines neuronal and synaptic preservation with reduced chronic glial activation and improved functional outcomes. They support further translational development of serotonin 1A receptor antagonism to limit secondary neurodegeneration and the visual and neurobehavioral consequences of TBI.

**Graphical Abstract:** 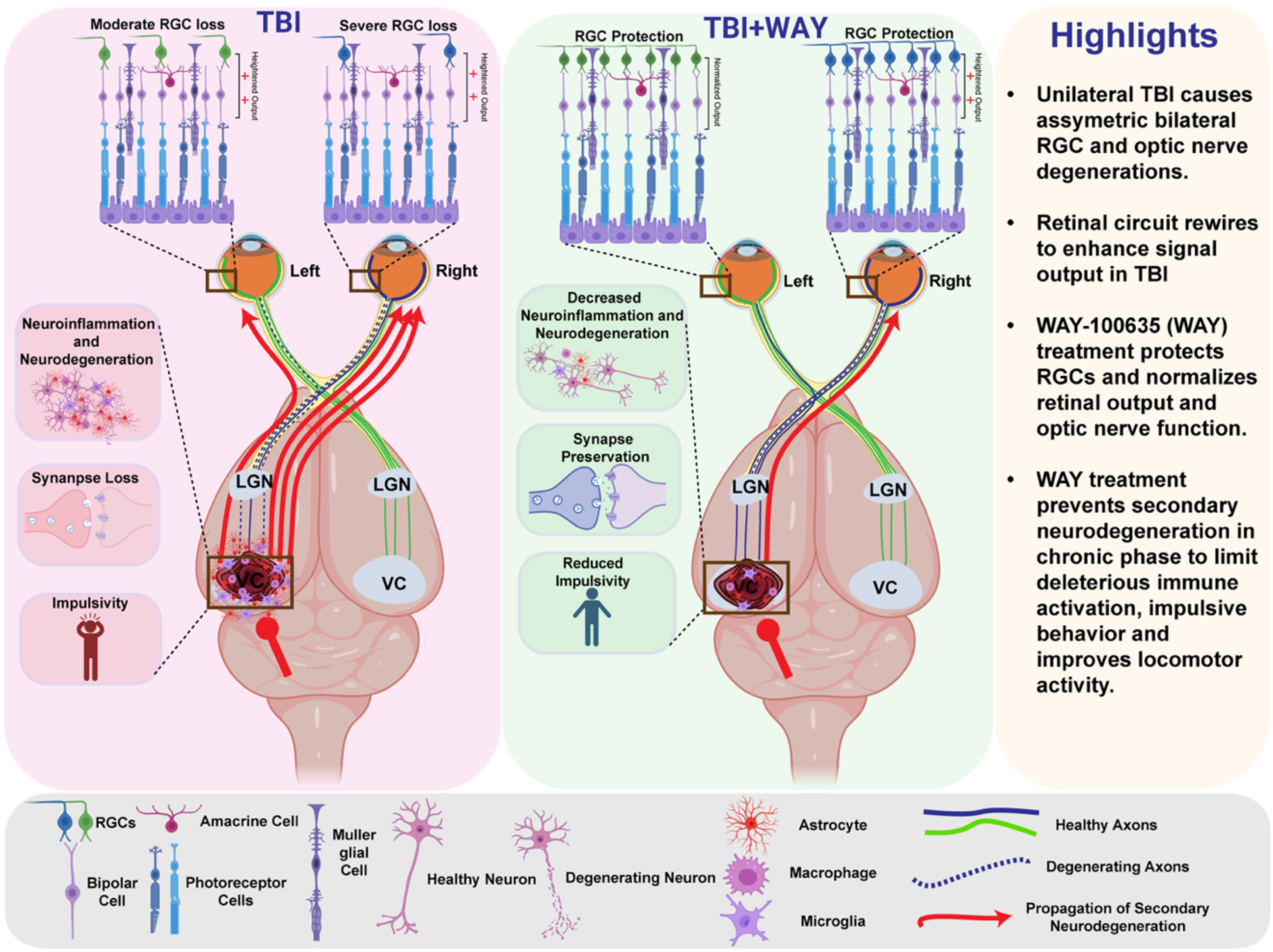

## Introduction

Traumatic brain injury (TBI) is a major health concern affecting service members, contact-sport athletes and civilians. TBI is increasingly recognized as the initiating event of a complex and progressive disease process encompassing visual dysfunction, neurodegeneration, anxiety and depressive disorders, and, in severe cases, increased risk of suicide^1,2^. Although TBI encompasses focal and diffuse injury, including cortical contusion and axonal damage, visual dysfunction is a frequent consequence of both concussion and more severe injury^3^. Visual complaints and oculomotor deficits are frequently reported among military personnel and veterans evaluated after TBI^3^. In a clinical cohort of 113 adolescents aged 11-17 years with persistent concussion symptoms, 70% had at least one oculomotor disorder when assessed 4-12 weeks after injury^4^.

Neurodegeneration after TBI evolves through overlapping acute and chronic phases. During the acute phase, the primary mechanical insult causes necrotic and lytic cell death, membrane disruption, and release of damage-associated molecular patterns (DAMPs), including intracellular proteins, nucleic acids, ATP, and mitochondrial constituents such as mitochondrial DNA. These signals activate pattern-recognition pathways in resident microglia and astrocytes and promote recruitment of peripheral myeloid cells, initiating a robust innate neuroimmune response^5,6^. As the injury progresses, incomplete recovery of inflammation, persistent mitochondrial dysfunctions, oxidative stress, impaired bioenergetics, and axonal damage sustain secondary neurodegeneration. Vulnerable neurons consequently undergo delayed regulated cell death, including apoptosis, in regions adjacent to and remote from the primary lesion, extending neuronal loss over days to weeks and contributing to chronic neurodegenerative pathology^7–9^.

Vision loss after TBI therefore emerges as part of a secondary neurodegenerative process that extends across the interconnected retino-visual pathway. RGCs are the projection neurons of the retina; their axons form the optic nerve and project to central visual targets, including the lateral geniculate nucleus (LGN), which relays retinal signals to the visual cortex (VC) through the optic radiations. Injury to the posterior visual pathway, including the visual cortex, can lead to retrograde trans-synaptic degeneration of RGCs^10,11^. Loss of target-derived trophic support may contribute to this process, although evidence that neurotrophins support RGC survival comes partly from developmental models and does not establish this mechanism after adult TBI^12,13^. Thus, TBI-induced degeneration of central visual neurons can potentially establish a feed-forward process in which mitochondrial dysfunction, axonal energetic failure, and loss of retrograde trophic signaling progressively destabilize the retina-to-brain circuit. Restoring mitochondrial homeostasis may therefore protect both RGCs and cerebral neurons vulnerable to secondary degeneration after TBI.

In support of TBI-associated visual circuit loss, experimental impact-acceleration injury induces multifocal axonopathy, neuroinflammation and RGC loss^14^, and blast injury produces early RGC axonal damage preceding subsequent soma loss^15^. Conceptually, this progression converges with glaucoma, although the initiating sites differ; following TBI, injury originates within the brain and can propagate retrogradely through the optic tract and optic nerve toward the retina, whereas glaucoma originates within the RGC-optic nerve projection and impairs neurotransmission and induces anterograde trans-synaptic degeneration of central visual targets^16^. Glaucoma models show early loss of RGC axonal transport cause distal axon degenerations, which recapitulate in patients with the evidence of optic nerve, LGN and visual cortex (VC) degenerations^17^. Thus, TBI and glaucoma may represent anatomically opposite but mechanistically convergent forms of degeneration across the same retino-visual axis, raising the possibility that an RGC-directed neuroprotective strategy effective in glaucoma may also protect interconnected visual pathway and brain neurons after TBI.

Traumatic optic neuropathy (TON) represents an important subset of TBI-associated visual pathology in which direct or indirect head trauma produces progressive optic nerve axon degeneration and RGC loss^18^. Corticosteroids and optic canal decompression have been used to treat TON^19^, but their efficacy remains uncertain. Separately, the Medical Research Council Corticosteroid Randomisation After Significant Head Injury (MRC CRASH) trial demonstrated increased mortality with high-dose corticosteroid treatment in patients with clinically significant head injury^20^. A major therapeutic gap remains; no established neuroprotective treatment prevents secondary degeneration associated with TON and TBI. Mitochondrial dysfunctions provide a mechanistic link between TON, RGC degeneration and cerebral TBI pathology. Secondary brain injury after TBI is characterized by inflammatory signaling-driven impaired mitochondrial bioenergetics, oxidative stress, and calcium dysregulation^21^. RGCs are particularly vulnerable to mitochondrial failure because their long partially myelinated axons require sustained energy production for electrical signaling and axonal transport^22^. These shared bioenergetic vulnerabilities support mitochondrial restoration as a therapeutic strategy for retinal and cerebral injury^23^.

We recently identified WAY-100635 (WAY) in a screen of human stem cell-derived retinal ganglion cells (hRGCs) for compounds that improve mitochondrial health. WAY protected hRGCs carrying a glaucoma-associated mutation and conferred neuroprotection in mouse models of optic nerve injury and glaucoma^22^. WAY is a well-characterized antagonist of the G protein-coupled 5-HT1A receptor^24^. Our previous experiments supported 5-HT1A specific signaling activation by WAY in hRGCs and demonstrated reversible activation of cyclic adenosine monophosphate (cAMP) signaling and PGC-1α-dependent mitochondrial biogenesis^22^.

Here, we asked whether WAY-100635, which we have discovered as a mitochondrial-restorative RGC-protective compound, could extend neuroprotection across the retina-brain axis following TBI. Using a controlled cortical impact model of TBI (CCI-TBI) involving the visual cortex, we found retinal and visual pathway dysfunctions, synaptic loss, and pronounced astrocytic and myeloid activation around the cortical injury site. These structural and functional pathologies were significantly attenuated by WAY along with improved early motor performance and attenuation of injury-associated increases in open-arm exploration as a readout for reduced anxiety like behavior. To our knowledge, this is the first evidence that a pharmacological compound that has been used in human^25,26^, preserves the retina-brain circuit and limits neuroinflammation and neurobehavioral dysfunction after TBI.

## Materials and methods

### Animals

All animal procedures were approved by the Institutional Animal Care and Use Committee (IACUC) at Indiana University School of Medicine and were conducted in accordance with the ARVO Statement for the Use of Animals in Ophthalmic and Vision Research and the ARRIVE guidelines. Wild-type male C57BL/6J mice (2-3 months old; The Jackson Laboratory) were randomly assigned to one of three groups [Sham (n = 3-10), TBI (n = 5-8), and TBI+WAY (n = 5-8)] following one week of acclimatization. Controlled cortical impact (CCI)-induced TBI was performed on day 0 in both the TBI and TBI+WAY groups. The TBI+WAY group received daily intraperitoneal injections of WAY-100635 (5 mg/kg body weight), beginning 1 hour after injury and continuing daily for 6 weeks until sacrifice, when eyes, optic nerves, and brains were collected for downstream analysis.

### Controlled cortical impact traumatic brain injury (CCI-TBI)

Mice were subjected to moderate CCI brain injury or sham surgery (craniotomy without impact) as previously described^27^, using an electromagnetic impactor (Impact One, Neuroscience Tools) targeting a site midway between lambda and bregma. Injury severity was standardized using a 3 mm tip diameter, 3.5 m/s contact velocity, 1.0 mm deformation, and 100 ms dwell time.

### Visual electrophysiology, behavioral vision testing, and diffusion MRI

RGC function was evaluated by flash ERG (Celeris system, Diagnosys) 31 days post-TBI using rod-saturating photopic flashes, with PhNR amplitude and implicit time quantified from the negative peak following the b-wave. Simultaneous fERGs and fVEPs were recorded 25-26 days after unilateral TBI; fERG waveforms were analyzed for b-wave and oscillatory potential (OP) amplitudes, and fVEP waveforms for peak amplitudes. Visual acuity and contrast sensitivity were assessed approximately 4 weeks post-TBI using the OptoMotry system (Cerebral Mechanics), based on head-tracking responses to drifting gratings. In vivo diffusion MRI was performed on a 9.4 Tesla Bruker system and data were fitted using diffusion tensor^28^, diffusion kurtosis^29^, and NODDI^30,31^ models, with probabilistic tractography (MRtrix3) used to assess fiber connectivity across the optic tract, LGN, and visual cortex^22,32,33^.

### Histology and immunofluorescence

Animals were transcardially perfused with 4% PFA, and brains and eyes were post-fixed, cryoprotected, and sectioned (30 μm) for immunohistochemical analysis. Retinas were harvested ∼6 weeks post-injury, immunolabeled for RBPMS, flat-mounted, and imaged by confocal microscopy; RBPMS-positive RGCs were quantified across four central and four peripheral regions per retina. Frozen coronal sections were stained with 0.1% cresyl violet for Nissl staining, and cortical cavity volume was quantified from these sections as the percentage difference between contralateral and ipsilateral spared cortex volume^34^. Sections were immunolabeled for NeuN, synaptophysin, GFAP, and CD68 as previously described^35^ and synaptophysin/GFAP intensity, NeuN-positive neurons, and CD68-positive cells were quantified in the perilesional cortex using ImageJ and an optical fractionator as previously described^36^.

### Behavioral testing

Motor coordination was assessed by rotarod (accelerating 0-30 rpm, 4 trials)^36^, and psychological-health-relevant risk-taking/behavioral disinhibition by EPM (10 min, ANY-maze tracking)^37^, quantifying time and distance in open versus closed arms.

### Statistics and reproducibility

Each animal was an independent biological replicate; sample sizes were based on group sizes used in comparable CCI-TBI studies ^36,38^. Group identity was masked, and optokinetic, electrophysiological, and dMRI acquisition/analysis were performed by experimenters blinded to treatment. One-way ANOVA with Tukey’s post-hoc correction was used for single-variable, three-group comparisons (Fig. 2B, C; Fig. 5E, I, M; Fig. 6K; Fig. 7B, D, E). Unpaired two-tailed Student’s t-tests were used where the question was restricted to two specific groups, including hemisphere- and eye-specific measures (Fig. 2F, G, I, J; Fig. 3B, C, E, F; Fig. 4D, E; Supplementary Fig. 1C, E; Supplementary Fig. 2C, E; Supplementary Table 1), and independently at each distance bin for GFAP intensity (Fig. 6L). Two-way ANOVA with Tukey’s post-hoc test assessed treatment versus eye effects (Fig. 3I). Analyses used GraphPad Prism 11.1; exact P values are reported in each figure, and data are mean ± SEM unless otherwise indicated.

Detailed methods are reported in the **Supplementary material**.

## Results

### Systemic WAY-100635 (WAY) treatment preserves the retino-visual circuit following TBI

We evaluated whether post-injury WAY treatment improves early functional recovery and protects against persistent secondary degeneration after CCI-TBI. TBI evolves from acute injury during the first few days through subacute recovery to chronic pathology characterized by persistent neuroinflammation and neurodegeneration^39^. Because CCI-induced motor deficits are greatest early and undergo substantial spontaneous recovery over 1-2 weeks^36^, rotarod performance was assessed at 3 days, 1 week and 2 weeks to capture acute impairment and subsequent recovery. Elevated-plus-maze (EPM) behavior was assessed at 2 weeks to examine neurobehavioral abnormalities beyond the immediate injury period^37^. Visual electrophysiology, optokinetic responses, diffusion MRI and histopathology were assessed at 3-6 weeks to determine whether WAY preserves neural structure and function during the chronic phase, beyond early motor recovery (**Fig. 1**).

**Figure 1.**
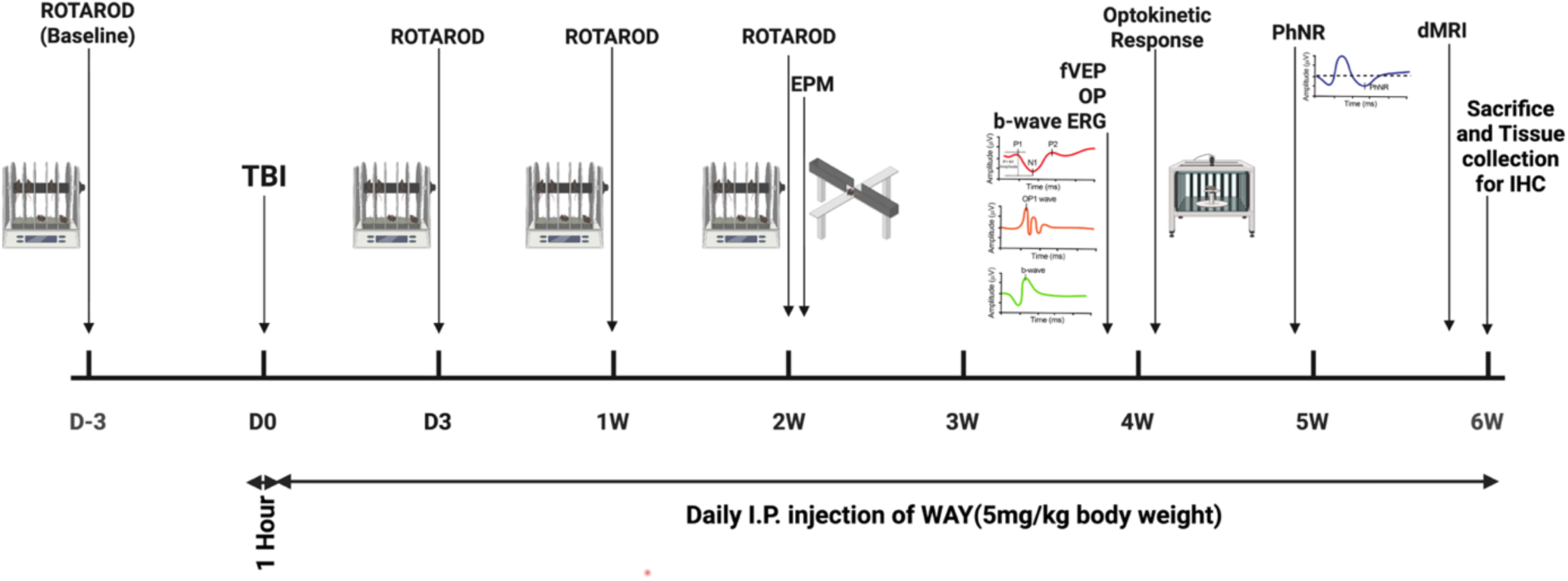
Experimental design and timeline of behavioral, electrophysiological, and imaging assessments after CCI-TBI. C57BL/6J mice underwent baseline rotarod testing prior to injury. Unilateral controlled cortical impact (CCI) TBI was induced at day 0, 1h after the injury daily intraperitoneal WAY-100635 injections (5 mg/kg body weight) continued until sacrifice. Motor function was assessed by rotarod at the acute and subacute phases at indicated timepoints following injury during which spontaneous recovery occurs. Elevated plus maze (EPM) testing for impulsive behavior was performed at indicated post-injury timepoints spanning subacute to chronic phases. Simultaneous flash visual evoked potential (fVEP), oscillatory potential (OP), and b-wave ERG recordings, along with optomotor response (OKR) testing for visual acuity and contrast sensitivity were obtained at subsequent indicated chronic phase timepoints. A photopic negative response (PhNR) recording was performed at chronic phase to assess inner retinal function. Diffusion MRI (dMRI) was acquired in the chronic phase shortly before sacrifice and tissue collection.

We assessed visual acuity (VA) and contrast sensitivity (CS) using the optokinetic response approximately 4 weeks after injury (day 31; **Fig. 2A**). CCI-TBI significantly reduced both measures compared with sham controls. Daily WAY treatment, initiated 1 hour after injury and continued throughout the study, significantly improved VA and CS compared with untreated TBI mice (**Fig. 2B, C**). These findings indicate preservation of behavioral visual function following post-injury treatment.

**Figure 2.**
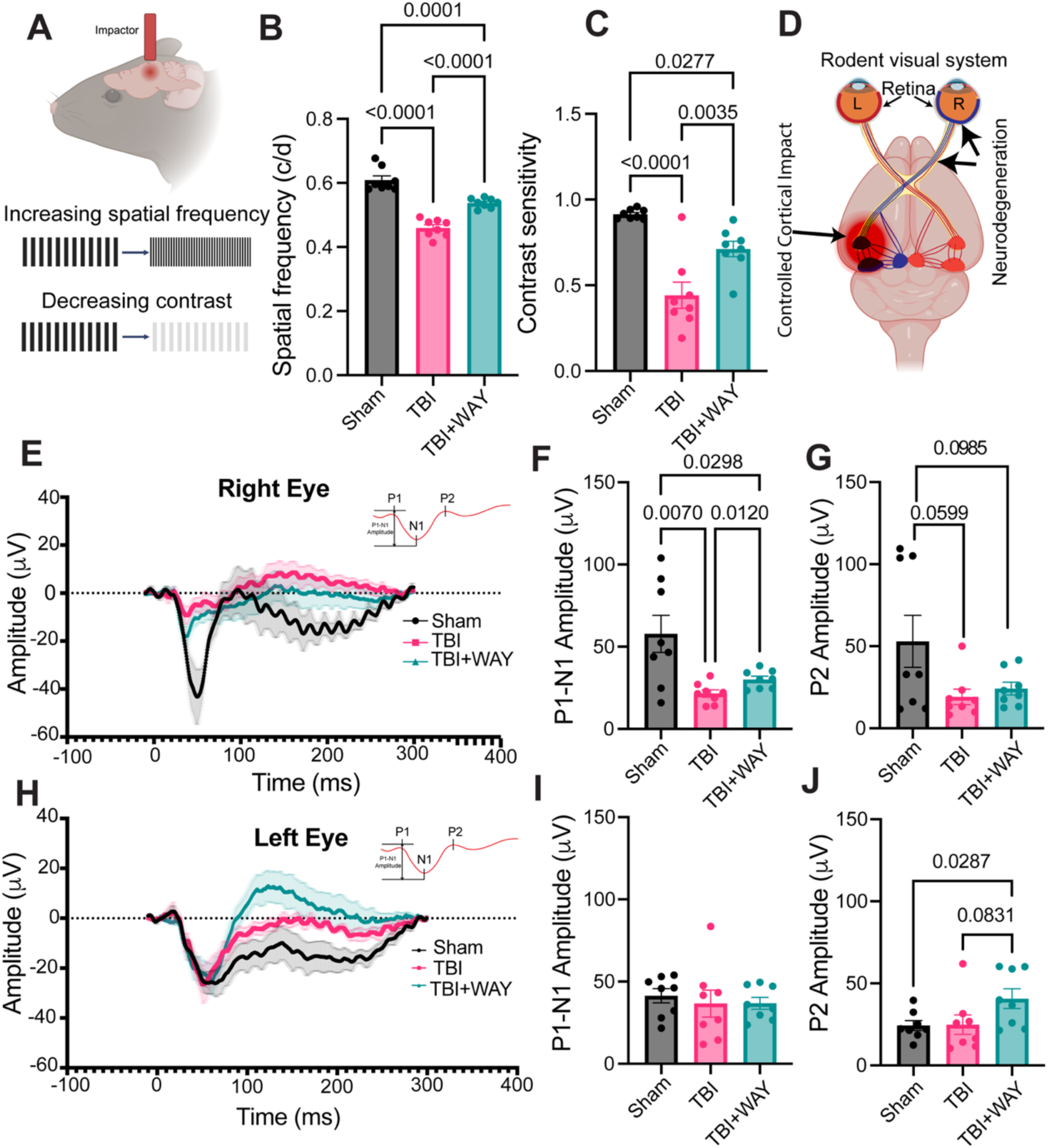
WAY treatment prevents retino-visual circuit integrity following CCI-TBI. **(A-C)** Functional visual outcomes assessed by optomotor response after around 4 weeks (Day 31) of injury. **(A)** Schematic illustration of the CCI-TBI model and visual assessment using the optomotor apparatus. **(B)** Quantification of Spatial Frequency (visual acuity, VA; cycles/degree) and, **(C)** Michelson contrast (contrast sensitivity, CS), determined by head-tracking responses to drifting gratings of increasing spatial frequency and decreasing contrast, respectively. Measurements were conducted under blinded conditions for both clockwise and counter-clockwise rotations. One-way ANOVA with Tukey’s multiple comparisons correction; *n* = 8 mice per group (Sham, TBI, TBI+WAY). Data are mean ± standard error of mean (SEM). **(D)** Schematic of murine visual projection anatomy illustrating the predominantly contralateral (∼90%) retinogeniculate pathway. **(E-J)** Flash visual evoked potential (fVEP) recordings after around 4 weeks (Day 25-26) of injury. **(E, H)** Representative fVEP traces from each experimental group; each trace represents the average per group. **(F, I)** Quantification of P1-N1 peak-to-peak amplitude (absolute values) under the indicated conditions. **(G, J)** Quantification of P2 peak amplitude under the indicated conditions. P2 amplitude reflects intracortical connectivity and is thought to index signal propagation from visual cortex to higher cortical areas. Statistical comparisons were done by an unpaired two-tailed t-test; *n =* 8 mice per group (Sham, TBI, TBI+WAY). Exact P values are indicated between compared groups in each panel.

Approximately 90% of mouse RGC axons cross at the optic chiasm^40^, so right-eye input predominantly reaches the injured left hemisphere (**Fig. 2D**). We assessed retino-cortical pathway function using fVEPs recorded simultaneously with fERGs at 25-26 days after injury. These cortical responses integrate retinal input and transmission through the central visual pathway^22,41^. Right-eye stimulation produced a significantly reduced P1-N1 amplitude in TBI mice, which was significantly improved by WAY treatment (**Fig. 2E, F**). P2 amplitude, reflecting later cortical visual processing^42^, remained significantly reduced for right eye in both untreated and WAY-treated TBI mice compared with sham; the treatment-associated improvement did not reach significance (**Fig. 2G**). Left-eye-evoked P1-N1 amplitude was comparable across groups (**Fig. 2H, I**), whereas P2 amplitude was significantly higher in WAY-treated mice than in both sham and untreated TBI mice (**Fig. 2J**). Together, these findings indicate that WAY preserves visually evoked responses in the pathway predominantly connected to the injured hemisphere, although recovery of later cortical responses remains incomplete. The increased P2 response to left-eye stimulation suggests improvement in lesion adjacent right cortical processing, presumably through a compensatory mechanism.

### WAY treatment preserves RGC structure-function and normalizes retinal interneuron circuitry following TBI

We assessed RGC function using the photopic negative response (PhNR), an RGC-weighted component of the photopic electroretinogram^43^, at 31 days after injury. TBI significantly reduced PhNR amplitude in both eyes compared with sham, indicating bilateral RGC dysfunction after unilateral left cortical injury (**Fig. 3A, B, D, E**). WAY treatment significantly preserved PhNR amplitude in both eyes (**Fig. 3B, E**). PhNR latency was significantly prolonged in the right eye after TBI and normalized by treatment (**Fig. 3C**), whereas no significant latency change occurred in the left eye (**Fig. 3F**). Together, these results indicate that WAY preserves bilateral RGC-weighted retinal output and prevents the right-eye delay associated with left cortical injury.

**Figure 3.**
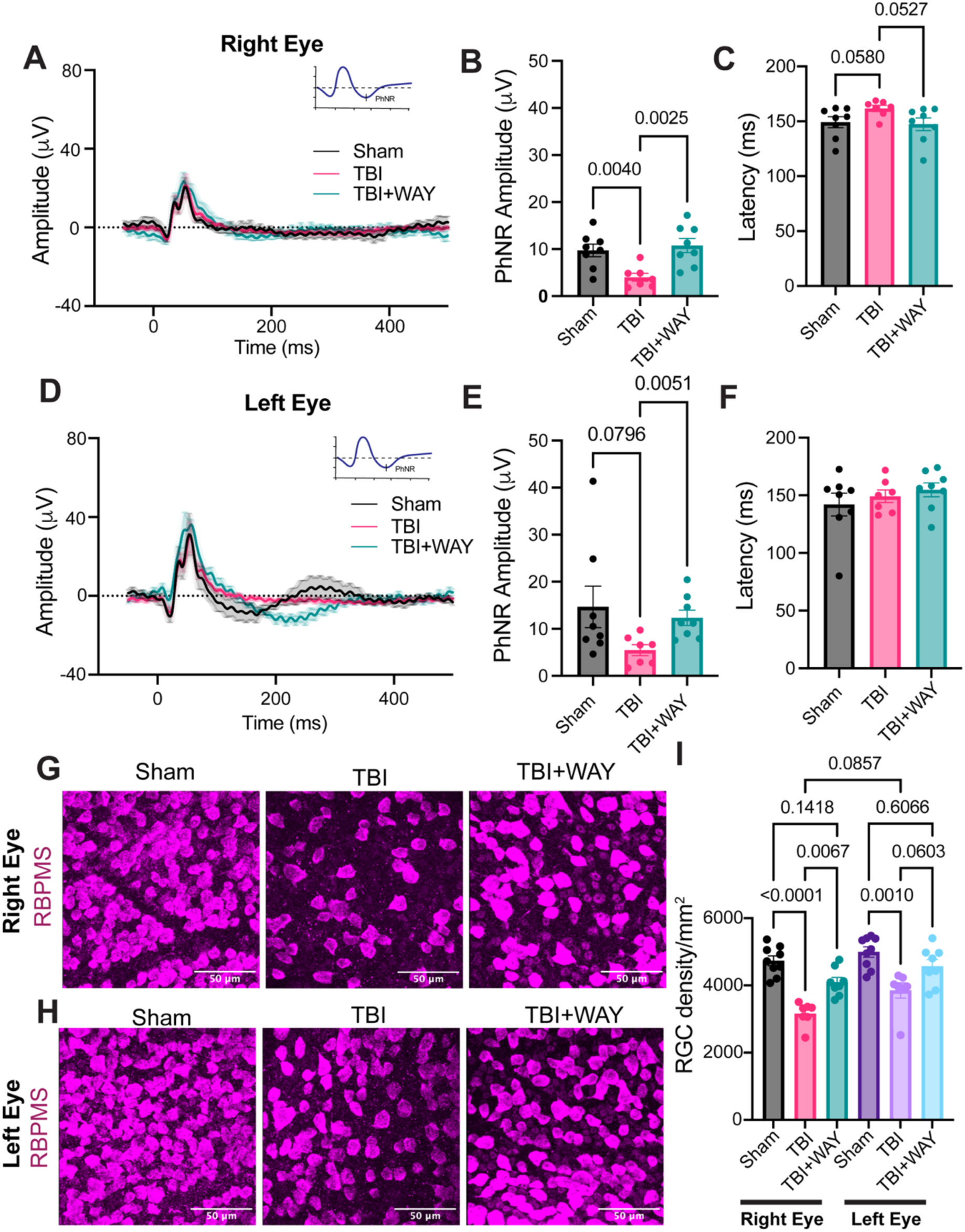
WAY treatment normalizes cone driven RGC response and prevents cell loss following CCI-TBI. **(A, D)** Representative flash electroretinogram (fERG) traces (white flashes on a rod-saturating green background) illustrating the photopic negative response (PhNR); each trace represents the average of all animals per group (three technical replicates per eye), recorded at approximately 5 weeks (Day 33) post-injury. **(B, E)** Quantification of PhNR amplitude (absolute values) under the indicated conditions. **(C, F)** Quantification of PhNR latency under the indicated conditions. Unpaired two-tailed t-test; *n =* 8 mice per group (Sham, TBI, TBI+WAY). Data are mean ± SEM. **(G, H)** Representative confocal images (40×/1.3 NA) of retinal whole-mounts immunolabeled for RBPMS, a marker of RGCs, were collected at approximately 6 weeks (Day 44) post-injury. **(I)** RGC density was quantified by counting RBPMS⁺ cells across four peripheral and four central regions per retina, averaged to yield a single value per eye. Data are mean ± SEM. Two-way ANOVA with Tukey’s post-hoc; n (Sham) = 9, n (TBI) = 7, n (TBI+WAY) = 8 mice per group. Exact P values are indicated between compared groups in each panel.

Because PhNR analysis revealed TBI-induced RGC dysfunction, we next examined whether inner-retinal circuitry was also altered. Oscillatory potentials (OPs) are high-frequency ERG components generated by synaptic interactions within the inner retina, particularly involving amacrine-mediated inhibitory networks (**Supplementary Fig. 1A**)^44,45^. TBI significantly increased OP1 amplitude in both eyes. In the left eye, WAY treatment reduced OP1 amplitude to sham levels, whereas the elevated OP1 response persisted in the right eye despite treatment (**Supplementary Fig. 1B-E)**, indicating differential rescue of inner-retinal circuit activity. In contrast, ERG b-wave amplitudes, which predominantly reflect photoreceptor-driven ON-bipolar cell activity^46^ (**Supplementary Fig. 2A)**, were largely preserved across groups in both eyes with a moderate reduction in the left eye of WAY-treated TBI mice (**Supplementary Fig. 2B-E)**. Together, these findings suggest that TBI preferentially disrupts inner-retinal neuronal circuitry rather than upstream photoreceptor-bipolar signaling, and that WAY partially normalizes this pathological retinal network response.

To determine whether functional preservation was accompanied by structural neuroprotection, we quantified RBPMS-positive RGCs in retinal flat mounts approximately 6 weeks after injury^47^. TBI significantly reduced RGC density in both eyes, with greater loss in the right eye. WAY-treated mice retained RGC densities comparable to sham in both eyes (**Fig. 3G-I**). These findings provide structural evidence of bilateral retinal neuroprotection accompanying the preservation of RGC function.

### Diffusion MRI reveals bilateral visual pathway abnormalities after TBI with localized WAY-mediated rescue

To assess retino-cortical microstructure, we acquired diffusion MRI (dMRI) and analyzed the data using three complementary models, alongside probabilistic tractography to reconstruct visual pathway trajectories. dMRI detects microstructural abnormalities that may be missed by conventional structural MRI^48^. Diffusion tensor imaging (DTI) measures diffusion directionality through fractional anisotropy (FA), and its magnitude through mean, axial and radial diffusivity (MD, AD and RD). Diffusion kurtosis imaging model (DKI) characterizes departures from Gaussian diffusion associated with tissue barriers and heterogeneity; mean, axial and radial kurtosis (MK, AK and RK) describe these departures overall and parallel or perpendicular to the principal diffusion direction, while kurtosis fractional anisotropy (KFA) measures their directional variation^49,50^. Neurite orientation dispersion and density imaging model (NODDI) estimates the intra-neurite fraction through neurite density index (NDI), angular dispersion through orientation dispersion index (ODI), and isotropically diffusing water through free-water fraction (FWF)^30,31,51,52^. These complementary measures provide a framework for interpreting visual pathway injury and treatment-associated changes^53^.

TBI to the left visual cortex was associated with bilateral microstructural abnormalities that extended bilaterally across the visual circuit (**Fig. 4A-C**; Supplementary Table 1). In the left LGN, radial kurtosis (RK) decreased significantly (Supplementary Table 1), indicating altered microstructural organization within this key relay of the retino-cortical visual pathway. The left OPT which is contiguous to the injured site, was more affected, with significant increases in FA, AD, KFA, AK, RK MK, and NDI (**Fig. 4D**). The concurrent increases in FA and NDI in the left OPT suggest altered microstructural organization, with greater directional diffusion coherence and increased neurite-like signal within the tract. At the chronic time point examined here, such changes should not be interpreted as improved tract integrity; rather, they may reflect reactive tissue compaction, cellular swelling, reduced extracellular water mobility, gliosis, or early remodeling before clear structural loss becomes dominant. This interpretation is consistent with reports that FA changes after TBI can be bidirectional and time-dependent^54,55^. Right-sided visual pathway involvement was also observed, anatomically consistent with the crossed organization of retinogeniculate projections at the optic chiasm^56^. Right OPT showed the most extensive diffusion changes, with significant increases in FA, AD, KFA, AK, RK, MK, and NDI, along with a significant decrease in FWF (**Fig. 4E**). The combination of increased FA and NDI may represent an apparent or injury-related increase in directional coherence and neurite-like signal, rather than a true gain in white matter integrity. Concurrent increases in kurtosis metrics, particularly MK and RK, further suggest increased diffusion restriction and microstructural heterogeneity, compatible with more severe tissue disruption within this tract.

**Figure 4.**
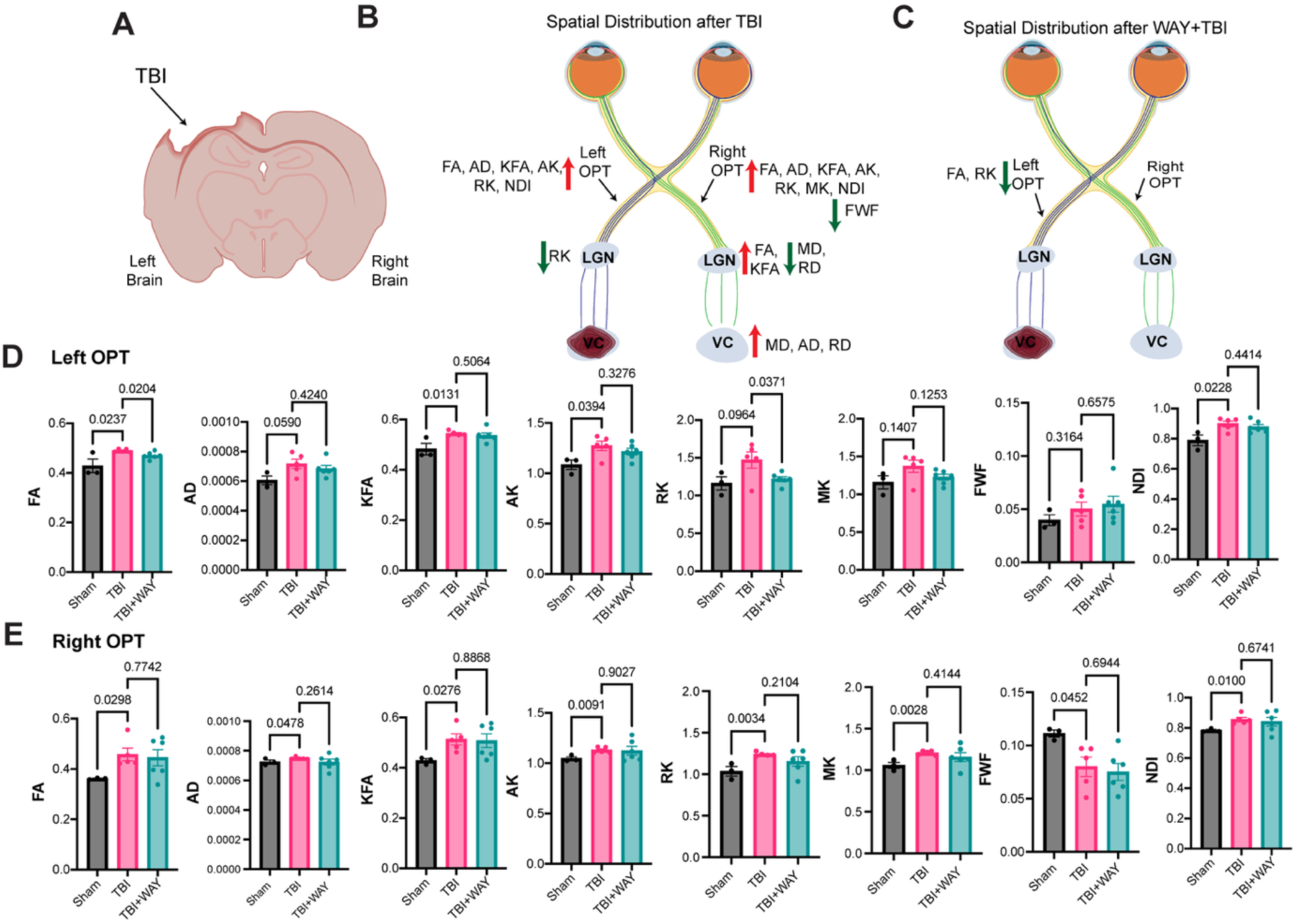
WAY treatment maintains microstructural organization of left optic tract following CCI-TBI. **(A)** Schematic of the unilateral CCI targeting the left visual cortex, examined around 6 weeks (Day 38-40) post-injury. (**B, C); (B)** Spatial maps of significant diffusion MRI alterations across the retino-cortical visual pathway during 38-40 days following TBI, **(C)** and after daily WAY treatment relative to untreated TBI. (**D, E); (D)** Quantification of diffusion metrics (FA, AD, KFA, AK, RK, MK, FWF, NDI) in the left OPT, (**E**) and right OPT across sham, TBI, and TBI+WAY groups. Unpaired two-tailed t-test; n (Sham) = 3, n (TBI) = 5, n (TBI+WAY) = 6 mice per group. Exact P values are indicated between compared groups in each panel. Data are mean ± SEM.

Because diffusion metrics are sensitive to tissue architecture but not specific to one pathological process, and FA reflects both microscopic anisotropy and orientation dispersion, these changes indicate altered tissue architecture rather than a single cell-type-specific process^57–59^. This is compatible with prior CCI-TBI studies showing complement activation, microglial remodeling, synaptic internalization and visual dysfunction in the dorsal LGN (dLGN) after unilateral cortical impact^60^.

The right visual cortex showed significant increases in MD, AD and RD (Supplementary Table 1), consistent with cortical tissue disruption and supporting a bilateral visual pathway response to unilateral cortical injury. This pattern is compatible with retrograde trans-synaptic degeneration after post-geniculate injury, as shown after restricted striate cortical lesions in macaques and acquired human occipital lobe, optic radiation and other retrogeniculate lesions^11,61,62^. The visual pathway is a recognized model of bidirectional trans-synaptic degeneration, in which pathology can spread across connected retinal, thalamic and cortical nodes in either direction depending on the site of the initiating lesion^63,64^.

In contrast to the widespread bilateral diffusion abnormalities after TBI, WAY treatment produced a highly localized neuroprotective effect within the left OPT, the region selected for mechanistic focus because it showed convergent multimodal diffusion abnormalities and significant WAY-mediated normalization (**Fig. 4D**). However, other left OPT metrics, including AK, MK, and NDI, were not significantly changed by WAY, although they trended toward improvement. Because AK and MK capture diffusion restriction along the principal axis and overall non-Gaussian microstructural complexity, their persistence suggests that internal axonal or broader neurite-associated abnormalities were only partially restored^29,49,50^. Thus, WAY appears to restore selected components of white matter microstructure, particularly RK and FA, without fully normalizing all diffusion signatures of axonal or neurite-associated injury. The selective reversal of these changes identifies the left OPT, adjacent to the injured left cortex as a key locus of WAY-mediated white matter neuroprotection after TBI.

### WAY treatment preserves synaptic integrity and reduces neuroinflammation following TBI

To evaluate cortical tissue loss and perilesional cellular pathology after TBI, we established a histological region-of-interest strategy centered on the injury site (**Fig. 5A**). CCI was targeted to include the left visual cortex. Coronal sections quantified gross cortical cavity formation, and sagittal sections assessed cellular and synaptic changes in spared perilesional cortex. To assess neuronal and synaptic preservation surrounding the lesion, neuronal nuclear antigen (NeuN), a marker of mature neurons^65,66^, was quantified within 0 - 600 μm of the cavity border. Synaptic integrity was evaluated by staining for synaptophysin, a well-established presynaptic vesicle protein^67,68^, within the more proximal 0 - 200 μm perilesional region as described in **Fig. 5A**.

**Figure 5.**
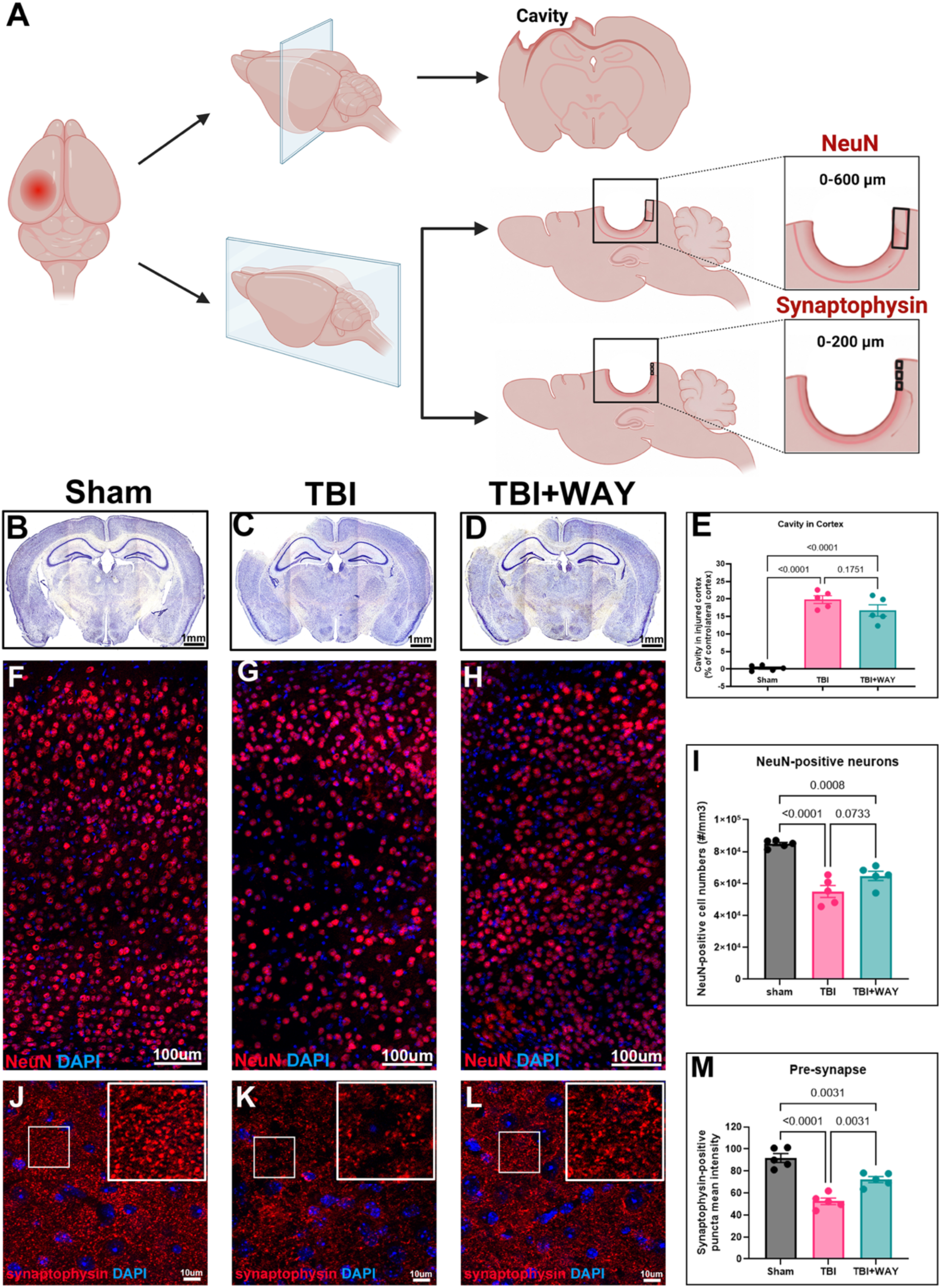
WAY treatment preserves synaptic integrity without affecting tissue loss following CCI-TBI. **(A)** Schematic representation of the coronal (separates front and back) and sagittal (separates left and right) brain section analyzed from the tissue collected approximately 6 weeks (Day 44) post-injury, showing the lesion cavity and the peri-lesional cortical regions selected for immunofluorescence imaging and quantitative analysis at defined distances from the cavity edge. **(B-D)** Representative Nissl-stained coronal brain sections from Sham, TBI, and TBI+WAY mice, respectively. **(E)** Quantification of cavity size in the ipsilateral injured cortex. **(F-H)** Representative images of NeuN immunostaining (red) showing surviving neurons in the spared visual cortex of Sham, TBI, and TBI+WAY mice, respectively. **(I)** Quantification of NeuN-positive cells in the spared visual cortex within a region 200-600 μm from the edge of the lesion cavity. **(J-L)** Representative images of synaptophysin immunostaining (red) in the spared visual cortex of Sham, TBI, and TBI+WAY mice, respectively. **(M)** Quantification of synaptophysin-positive fluorescence intensity in the spared visual cortex, demonstrating significantly greater synaptic preservation following WAY treatment after TBI. Data are mean ± SEM. One-way ANOVA with Tukey’s post-hoc; n (Sham, TBI, TBI+WAY) = 5. Exact P values are indicated between compared groups in each panel.

To test whether WAY preserves brain tissue after TBI, Nissl staining of coronal sections at the injury epicenter showed that sham surgery left ipsilateral morphology unchanged relative to the contralateral hemisphere (**Fig. 5B**). In contrast, TBI produced a large cortical cavity, consistent with the focal tissue loss and progressive cavitation characteristic of CCI injury (**Fig. 5C**)^27,69,70^. Although the lesion cavity appeared smaller in WAY-treated mice (**Fig. 5D**), quantitative analysis of lesion volume showed that this reduction did not reach statistical significance (**Fig. 5E**).

Using the mouse brain atlas as a reference, we found that our CCI model damaged approximately one-third of the visual cortex. Because a substantial portion of the cortex remained intact, we next asked whether neurons within this spared region were also affected by the injury and if WAY treatment protected these neurons. NeuN-positive neurons were stereologically quantified in the peri-lesional visual cortex (0 - 600 μm from the lesion border). As expected, TBI significantly reduced the number of surviving neurons compared with sham controls (**Fig. 5F, G and I**). WAY-treated mice showed a significantly higher number of NeuN-positive neurons than untreated TBI mice, supporting a neuroprotective effect (**Fig. 5G-I**).

While neuronal survival showed an improvement, synaptic preservation was much more pronounced. Synaptophysin immunoreactivity was substantially higher in the spared visual cortex of WAY-treated mice than in untreated TBI animals (**Fig. 5J, K and L**). This observation was supported by quantitative analysis of fluorescence intensity, which demonstrated significant preservation of synapses following WAY treatment (**Fig. 5M**).

Neuroinflammation is a major contributor to synaptic loss and secondary neurodegeneration after TBI^71^. Because WAY markedly preserved synapses, we next investigated whether this protection was accompanied by reduced neuroinflammation (**Fig. 6A**). WAY treatment dramatically reduced CD68 immunoreactivity (**Fig. 6B-D, H-J, K),** a marker for activated macrophages and phagocytic microglia^72^, and significantly attenuated GFAP expression, reflecting reduced reactive astrogliosis^73^, compared to untreated TBI animals (**Fig. 6E-J, L**). These findings indicate that WAY suppresses both microglial and astrocytic neuroinflammatory responses in the perilesional cortex.

**Figure 6.**
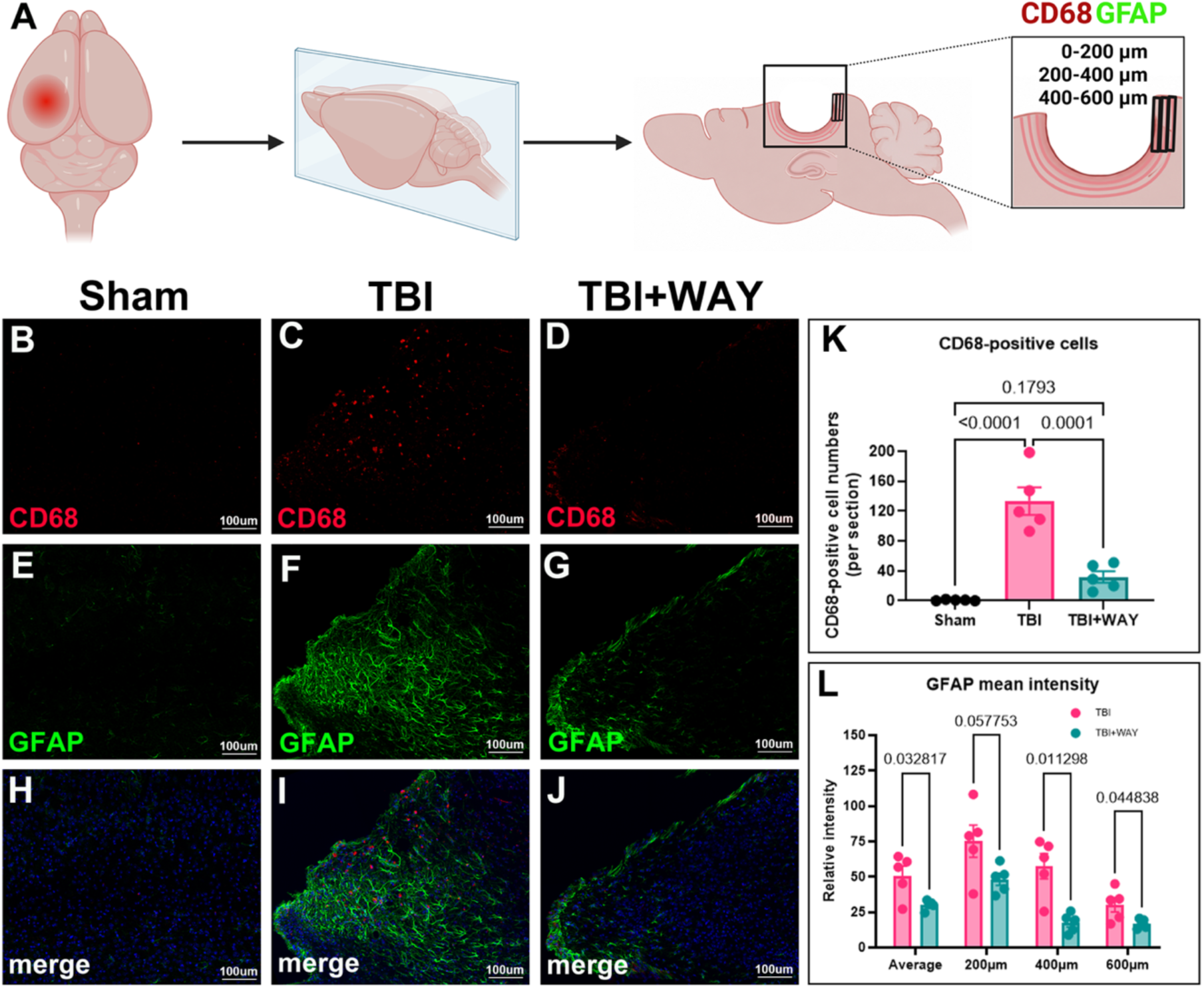
WAY treatment attenuated neuroinflammation response in the injured brain lesion following CCI-TBI. **(A)** Schematic representation of the sagittal brain section analyzed from the tissue collected approximately 6 weeks (Day 44) post-injury, showing the lesion cavity and the peri-lesional cortical regions selected for immunofluorescence imaging and quantitative analysis at defined distances from the cavity edge. **(B-J)** Representative immunofluorescence images showing CD68 (red) and GFAP (green) immunoreactivity in the injured and peri-lesional cortex of Sham, TBI, and TBI+WAY mice. **(B, E)** Minimal CD68-positive signals and low GFAP immunoreactivity were detected in the cortex of Sham mice, respectively; **(H)** merged image of **B**, and **E**. **(C, F)** TBI induced robust increases in CD68 and GFAP immunoreactivity at the lesion site and surrounding cortical regions, respectively; **(I)** merged image of **C** and **F**. **(D, G)** WAY treatment significantly reduced TBI-induced CD68 and GFAP immunoreactivity, respectively; (**J)** merged image of **D** and **G**. **(K)** Quantification of CD68-positive cells in the cortex of Sham, TBI, and TBI+WAY mice. **(L)** Quantification of GFAP fluorescence intensity at defined distances from the edge of the lesion cavity. Data are mean ± SEM. One-way ANOVA with Tukey’s post-hoc for **K** and multiple unpaired t-tests for **L**; n (Sham, TBI, TBI+WAY) = 5. Exact P values are indicated between compared groups in each panel.

### WAY treatment improves neuropsychiatric behavior following TBI

The preservation of cortical neuronal and synaptic markers, together with reduced glial activation, prompted us to assess functional effects beyond vision. We evaluated motor coordination using the rotarod and exploratory behavior using the elevated plus maze (EPM) which is an established assay for psychological-health-relevant risk-taking/behavioral disinhibition (**Fig. 7A, C**)^36,74,75^.

**Figure 7.**
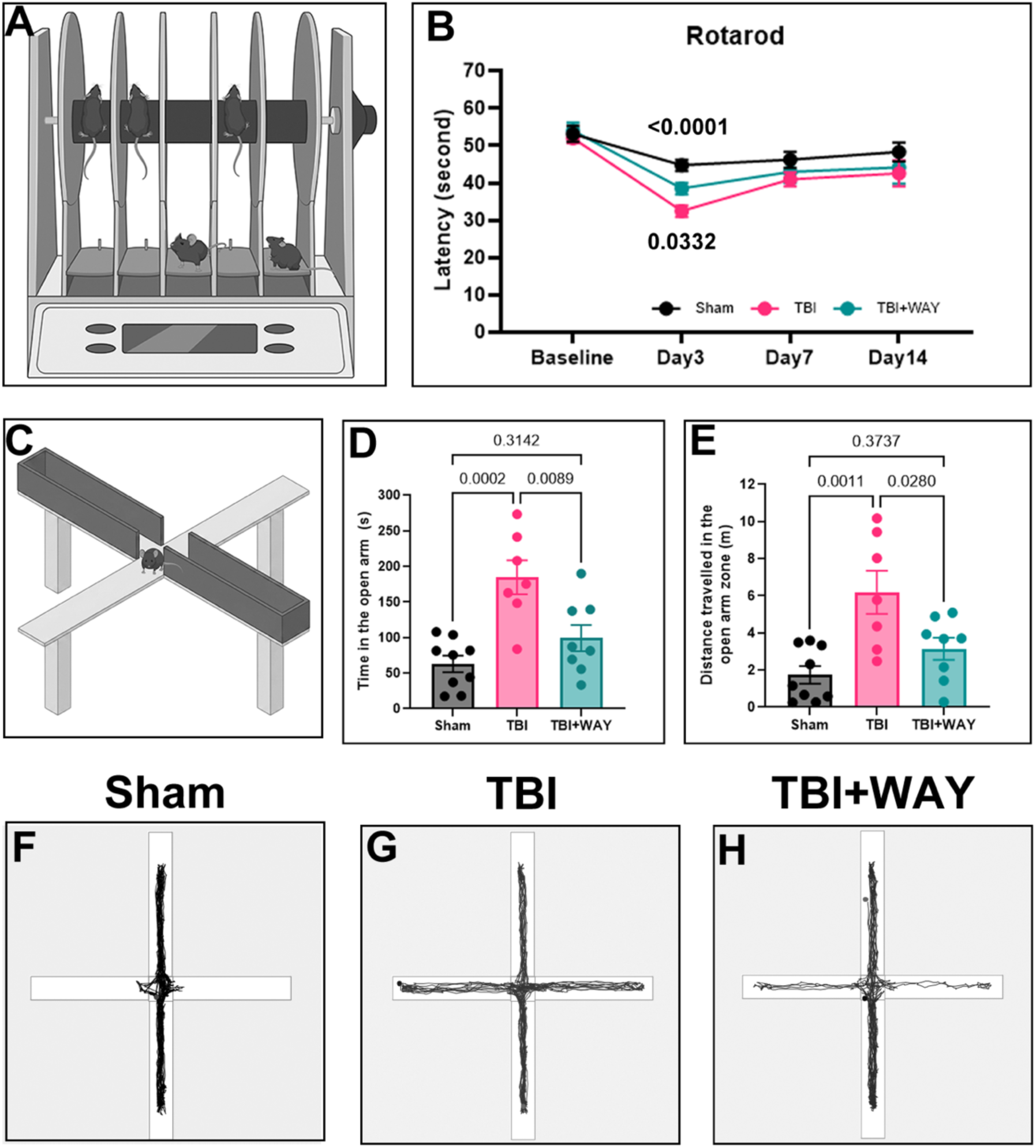
WAY treatment improved motor and emotional function outcomes following CCI-TBI. **(A)** Schematic representation of the of rotarod apparatus. **(B)** Quantification of latency to fall from the rotating rod on days −3, 3, 7 and 14 after TBI. Data are mean ± SEM. One-way ANOVA with Tukey’s post-hoc; n (Sham) = 10, n (TBI, TBI+WAY) = 8 mice per group. Exact P values are indicated between compared groups in each panel. **(C)** Schematic representation of the elevated plus maze (EPM) apparatus. **(D, E)** Quantification of open-arm exploration in the elevated plus maze 2 weeks after TBI, measured as time spent in the open arms and distance traveled within the open arms. **(F-H)** representative tracking plots from sham, TBI, and TBI+WAY mice during EPM test, respectively. Data are mean ± SEM. One-way ANOVA with Tukey’s post-hoc; n (Sham) = 9, n (TBI) = 7, n (TBI+WAY) = 8 mice per group. Exact P values are indicated between compared groups in each panel.

TBI significantly impaired motor performance, as reflected by a shorter latency to fall compared with sham-operated mice (**Fig. 7B**). Mice treated with WAY generally performed better than untreated TBI mice throughout the testing period, with the most noticeable improvement occurring during the acute phase of recovery. On day 3 after injury, WAY-treated mice remained on the rotating rod significantly longer than untreated TBI mice (**Fig. 7B**).

The EPM revealed another important consequence of TBI. Compared with sham animals, TBI mice spent more time in the open arms and traveled a greater distance within the open arms, indicating psychological-health-relevant risk-taking/behavioral disinhibition (**Fig. 7D-G**). This phenomenon aligns with prior CCI studies reporting increased open-arm exploration in the EPM and with reviews noting that TBI-related increases in open-arm activity may reflect impulsivity, altered arousal, or disinhibition^75^. These behavioral abnormalities were significantly attenuated by WAY treatment, as treated mice spent less time and traveled a shorter distance in the open arms than untreated TBI mice (**Fig. 7D-H**).

## Discussion

TBI remains without an established pharmacological therapy that prevents secondary neurodegeneration, and treatment options are particularly limited for TBI-associated visual pathway injury and traumatic optic neuropathy (TON). This therapeutic gap is important because TBI is not restricted to the primary cortical lesion but initiates progressive mitochondrial dysfunction, axonal injury and neuroinflammation that can compromise interconnected neural systems long after the initial trauma^5–9^. Our study identifies the human tested 5-HT1A antagonist WAY-100635 (WAY) as a post-injury neuroprotective strategy with efficacy extending from the retina and optic pathway to the injured brain. The convergence of electrophysiological, behavioral, diffusion MRI and histological outcomes argues that WAY does not simply improve a single TBI phenotype but limits secondary degeneration across anatomically connected neural compartments. This extends our previous demonstration of WAY-mediated mitochondrial restoration and RGC protection in optic nerve injury and glaucoma^22^, and provides a strong foundation for defining therapeutic window, dosing, pharmacokinetics and safety required for translational development.

An important finding was the asymmetric relationship between retinal and cortical electrophysiology after unilateral left CCI. Approximately 90% of mouse RGC projections cross at the optic chiasm^40^; therefore, right-eye input predominantly enters the injured left visual pathway, whereas most left-eye axons project toward the relatively spared right hemisphere. This organization explains the pronounced right-eye fVEP deficit, but PhNR and RGC count measurements nevertheless revealed RGC dysfunction and degeneration in both eyes. These observations are not contradictory because fVEP and PhNR interrogate different compartments; PhNR directly reflects RGC-weighted inner-retinal function^43,76^, whereas fVEP requires sufficient retinal input, axonal transmission and cortical recruitment^41,42^. Partial RGC loss in the left eye may therefore be detectable by PhNR without reducing input below the threshold needed to compromise the cortical fVEP response. Moreover, posterior visual pathway lesions can produce retrograde trans-synaptic RGC degeneration, including after striate cortical injury and head trauma^11,61–64,77^. The preferential right-eye PhNR latency abnormality and greater right-eye RGC loss further supports a stronger secondary degeneration along the projection directly coupled to the injured hemisphere.

The oscillatory potential findings indicate that this secondary retinal pathology extends beyond RGCs to inner-retinal network function. Oscillatory potentials arise largely from synaptic interactions involving amacrine and other inner-retinal neurons^44,45^, whereas preservation of the b-wave argues against comparable disruption of upstream photoreceptor-ON-bipolar signaling. The bilateral OP1 elevation after TBI therefore suggests maladaptive remodeling or hyperactivity within inner-retinal circuitry. WAY normalized this response in the left eye but not the more strongly affected right-eye pathway. The right retina, whose predominant projection traverses the injured hemisphere, may undergo greater secondary circuit remodeling that persists despite largely preserved RGC survival and output. In contrast, the less severely affected left retinal network may remain more functionally reversible. Because oscillatory potentials integrate activity across several inner-retinal cell populations, these findings should not be interpreted as selective amacrine-cell protection; rather, they demonstrate that WAY differentially preserves retinal network homeostasis according to the severity and anatomical relationship of each projection to the brain lesion.

Diffusion MRI provides an independent view of this anatomically heterogeneous injury. Within the left OPT, WAY reduced the elevated RK value observed in TBI animals and normalized FA toward baseline. The increased RK following TBI indicates altered non-Gaussian diffusion perpendicular to the principal tract orientation, potentially reflecting changes in tissue heterogeneity and compartmental water distribution. Brain extracellular diffusion is strongly shaped by extracellular space (ECS) volume fraction and tortuosity^78^, while TBI-associated cytotoxic/cellular edema involves water shifts into cells and reduced diffusion mobility^79,80^. Under these conditions, increased FA and NDI should not be interpreted as improved white matter integrity; rather, they may represent an apparent increase in directional coherence and neurite-like signal caused by restricted radial diffusion and altered compartmental water distribution^30,31,54,57,81^. This interpretation is further supported by experimental TBI studies showing that increased anisotropy can reflect reactive gliosis rather than axonal regeneration or true structural improvement^82^. Thus, WAY-mediated reduction of RK and normalization of FA support attenuation of selected microstructural abnormalities, although their cellular substrates remain unresolved.

The cortical histology further distinguishes irreversible primary damage from therapeutically accessible secondary injury. CCI produces immediate mechanical tissue destruction followed by progressive cavitation. The preservation of perilesional neuronal and synaptic markers without a significant reduction in cavity volume is consistent with protection of surviving tissue vulnerable to secondary injury. In contrast, neurons and synapses surrounding the lesion remain exposed to delayed excitotoxic, metabolic and inflammatory stress and are potentially salvageable. Preservation of NeuN-positive neurons and particularly synaptophysin-positive terminals by WAY is therefore more consistent with inhibition of secondary degeneration than repair of the primary lesion. This distinction is important because synaptic dysfunction can precede neuronal loss and may represent one of the earliest reversible substrates of neurological impairment after TBI.

Microgliosis and astrogliosis have been observed up to a year after a single TBI^83^. The marked reduction of CD68-positive phagocytic microglia/macrophages and GFAP-positive reactive astrocytes in the chronic phase provides further evidence that WAY modifies the chronic secondary-injury environment. Persistent neuronal degeneration releases DAMPs that sustain glial activation, inflammatory signaling and blood-brain barrier (BBB) dysfunction well beyond the acute injury period^5,6,84^. BBB disruption can, in turn, facilitate peripheral immune-cell recruitment and amplify neuroinflammation, generating a self-reinforcing cycle of neuronal injury and inflammatory activation^84^. Thus, the coordinated preservation of neurons and synapses together with reduced CD68 and GFAP responses is consistent with interruption of this chronic neurodegenerative-neuroimmune feedback loop. Reduced CD68 may reflect diminished macrophage recruitment and improved BBB integrity, but CD68 labels both resident microglia and infiltrating macrophages, so BBB protection is not directly shown; vascular permeability and tight-junction assays are needed. Nonetheless, the chronic suppression of both microglial/macrophage and astrocytic responses strongly supports a neuroprotective environment generated by WAY rather than merely an effect on acute lesion formation.

Retinal and brain protection may reinforce each other through the reciprocal trophic organization of the visual pathway; central targets supply RGCs with retrogradely transported BDNF and neurotrophin-4/5, and their disruption accelerates RGC death^12,13^, while preserved RGC axons sustain visual input and trophic support to the LGN, limiting deafferentation. TBI can therefore initiate a deleterious feed-forward loop in which central visual pathway degeneration reduces retrograde support to RGCs, while declining RGC input further destabilizes central targets. By preserving RGC structure and function, WAY may interrupt the retinal component of this loop. Because WAY was administered systemically and also preserved perilesional neurons and synapses, it may simultaneously protect cerebral neurons from secondary metabolic stress. Such reciprocal protection provides a mechanistic framework by which maintaining both ends of the retina-brain circuit could progressively stabilize the entire pathway.

The behavioral effects of CCI further emphasize that focal cortical trauma evolves into a distributed brain disorder. Although the impact was centered around visual cortex, secondary injury after CCI affects connected thalamic and cortical networks and can produce persistent neurobehavioral abnormalities^83,85^. Motor deficits after experimental CCI are typically greatest during the early post-injury period and undergo substantial spontaneous recovery over the following days to weeks^36^. The greater early rotarod recovery with WAY therefore suggests acceleration of functional restoration during a period when injured but viable networks remain capable of compensation. Likewise, altered elevated-plus-maze (EPM) behavior after CCI has been reported across injury severities and can reflect altered anxiety, threat assessment, impulsivity or disinhibition rather than a unitary anxiety phenotype. The normalization of these behaviors by WAY, together with reduced cortical neuroinflammation and preserved synaptic connectivity, suggests that its neuroprotective actions extend beyond the visual pathway to distributed neural networks governing motor and affective behavior.

Previous studies of 5-HT1A receptor targeting after TBI have primarily investigated agonist treatment. A single dose of 8-OH-DPAT, which activates both 5-HT1A and 5-HT7 receptors, improved behavioral recovery when administered 15 minutes after experimental injury but was ineffective when delayed to 1 or 2 hours^86^. Subsequent experiments showed that 5-HT7 blockade with SB-269970, but not 5-HT1A blockade with WAY-100635, abolished its cognitive benefit, implicating 5-HT7 signaling in that treatment paradigm^87^. The selective 5-HT1A agonist repinotan was also evaluated in a small safety and tolerability trial in severe TBI, but therapeutic efficacy was not established^88^. Our study extends this literature by identifying repeated post-injury WAY-100635 treatment as an antagonist-based approach with protective effects evident during the chronic phase, broadening the therapeutic rationale beyond the acute agonist paradigms described above.

Mechanistically, these findings are consistent with our previous demonstration that WAY restores mitochondrial homeostasis in stressed RGCs through 5-HT1A-dependent signaling and cAMP-PGC-1α-mediated mitochondrial biogenesis^22^. Mitochondrial bioenergetic failure and neuroinflammation are tightly coupled components of secondary TBI pathology. Mitochondrial dysfunction increases oxidative and inflammatory stress, while sustained inflammatory signaling further compromises oxidative phosphorylation and neuronal viability. The present study does not establish that mitochondrial restoration alone accounts for the cerebral effects of WAY, but the convergence of preserved neurons and synapses, reduced chronic glial activation, improved white-matter microstructure and functional recovery is consistent with interruption of this metabolic-inflammatory cycle. Defining whether WAY acts directly on neuronal 5-HT1A signaling, neuroimmune cells, or both will be an important direction for future studies.

Our findings support a model in which WAY-100635 protects the vulnerable neural circuitry surrounding and connected to it from progressive secondary degeneration. By preserving RGCs and visual signaling, limiting optic pathway microstructural disruption, maintaining cortical neurons and synapses, suppressing chronic neuroimmune activation, and improving neurological and behavioral outcomes, WAY provides broad protection across the retina-brain axis. Together with prior human PET studies demonstrating brain accessibility of radiolabeled WAY-100635 at tracer exposure^25^, these findings support further translational development, including evaluation of exposure-response relationships and safety at therapeutic doses.

## DECLARATIONS

### Availability of data and materials

The data supporting the findings of this study are available within the article and its Supplementary Information. Quantitative dMRI metrics and the corresponding statistical analyses are provided in Supplementary Table 1. Additional data and resources generated or analyzed during the current study are available from the corresponding author upon reasonable request. Requests for further information, data, and resources should be directed to the lead contact, Arupratan Das.

