## Supplementary Information for "5-HT1A Antagonist WAY-100635 Prevents Chronic Neurodegeneration and Immune Activation After TBI"

**Supplementary Methods**

**Animals**

All animal procedures were approved by the Institutional Animal Care and Use Committee (IACUC) at Indiana University School of Medicine and were conducted in accordance with the ARVO Statement for the Use of Animals in Ophthalmic and Vision Research and the ARRIVE guidelines. Wild-type male C57BL/6J mice (2-3 months old) were obtained from The Jackson Laboratory (Bar Harbor, ME, USA) and maintained under standard housing conditions at the Laboratory Animal Resource Center (LARC), Indiana University School of Medicine.

For the controlled cortical impact (CCI) model, mice were randomly assigned to one of three groups [Sham (n = 3-10), TBI (n = 5-8), and TBI+WAY (n = 5-8)] following one week of acclimatization. CCI-induced TBI was performed on day 0 in both the TBI and TBI+WAY groups. The TBI+WAY group received daily intraperitoneal injections of WAY-100635 (5 mg/kg body weight), beginning 1 hour after injury and continuing daily for 6 weeks until sacrifice. At the experimental endpoint, animals were euthanized, and eyes, optic nerves, and brains were collected for downstream analysis.

**Controlled cortical impact traumatic brain injury (CCI-TBI)**

Mice (2-3 months old) were subjected to moderate, controlled cortical impact (CCI) brain injury or sham surgery as previously described(*1*). Briefly, the mice were anesthetized and placed in a stereotaxic frame (Kopf Instruments, Tujunga, CA). Using sterile procedures, the skin was retracted, and a 4 mm craniotomy was performed. A point was identified midway between the lambda and bregma sutures and laterally midway between the central suture and the temporalis muscle. The skullcap was carefully removed without disruption of the underlying dura by drilling. Prior to injury, the tip of the impactor was angled and kept perpendicular to the exposed cortical surface. The mouse CCI model used an electromagnetic impactor (impact one, Neuroscience Tools, O’Fallon, MO) that allowed to adjust the severity of the injury by tip diameter, controlling contact velocity, the level of cortical deformation, and the impact dwell time independently. To generate brain injury with moderate severity, the tip with 3mm diameter was selected, contact velocity was set at 3.5 m/s, deformation at 1.0 mm, and dwell time at 100 ms. During surgery and recovery, the core body temperature of the animals was maintained at 36-37 °C using a heating pad. Sham animals were subjected to craniotomy but did not receive an impact.

**Photopic negative response (PhNR) recording by flash electroretinogram (fERG)**

RGC function was evaluated using the Celeris system (Diagnosys, Inc., Lowell, MA) paired with Espion software (Diagnosys, Inc.). ERG recordings were obtained 31 days post-TBI. Prior to testing, mice underwent overnight dark adaptation, and all subsequent procedures were conducted under dim red light (660 nm). Animals were anesthetized with isoflurane and placed on a temperature-controlled platform maintained at 37 °C. Pupil dilation was achieved with 1% tropicamide (Somerset Therapeutics, FL), and topical anesthesia was administered via 0.5% proparacaine hydrochloride (Bausch + Lomb, NJ). Corneal moisture was preserved throughout using Systane lubricant (Alcon, TX).

A contact lens-style probe incorporating an integrated electrode, stimulator, and pinhole aperture was positioned on the cornea and centered over the pupil. The reference electrode was secured on the contralateral eye. PhNR responses were elicited through 100 consecutive white-light flashes (20.0 cd·s/m²) delivered against a rod-saturating green background (40 cd·s/m²) to provoke RGC-specific activity. Raw signals underwent bandpass filtering at 0.125-40 Hz to reduce oscillatory artifacts. To ensure recording consistency, the probe was repositioned at a minimum of three distinct angles per animal, and the resulting PhNR amplitudes were averaged. PhNR amplitude was defined as the voltage difference between baseline and the negative peak immediately following the b-wave; implicit time (latency), defined as the interval from stimulus onset to that peak, was recorded concurrently. Following completion of the procedure, animals were returned to their home cages and placed on heating pads covered with drapes. Cages were arranged so that only half of each cage rested on the heating pad (low temperature setting), enabling mice to self-regulate their body temperature. Animals were monitored until fully ambulatory (approximately 10-15 minutes) before being returned to the animal facility.

**Simultaneous flash ERG and flash visual evoked potential (fVEP) recording**

Simultaneous flash electroretinograms (fERGs) and flash visual evoked potentials (fVEPs) were recorded using the Celeris system integrated with Espion software (Diagnosys, Lowell, MA), 25-26 days following unilateral TBI. Animals underwent overnight dark adaptation prior to testing, and all handling was performed under dim red light (660 nm). Mice were anesthetized with ketamine/xylazine (90/10 mg/kg, i.p.) and placed on a temperature-controlled platform maintained at 37 °C. Pupils were dilated with 1% tropicamide (Somerset Therapeutics, FL), and topical corneal anesthesia was achieved with 0.5% proparacaine hydrochloride (Bausch + Lomb, NJ). Corneal hydration was maintained with PBS throughout the recording session.

A contact lens electrode incorporating an integrated stimulator and pinhole aperture was centered over the pupil of the stimulated eye. The reference electrode was placed on the contralateral eye, subdermal platinum needle electrodes were inserted into the cheek and over the occipital scalp overlying visual cortex, and the ground electrode was positioned at the base of the tail. Each eye was stimulated and recorded independently. Simultaneous fERG and fVEP responses were elicited using 50 white-light flashes (6500 K, 1 cd·s/m²). Flash ERG waveforms were analyzed for b-wave amplitude and oscillatory potential (OP) amplitudes; fVEP waveforms were analyzed for peak amplitudes. All measurements were compared across groups using GraphPad Prism.

**Optokinetic response (OKR) testing**

Visual function was evaluated approximately 4 weeks after unilateral TBI (Day 31) using the OptoMotry system (Cerebral Mechanics, Inc.). Optokinetic response (OKR) testing was performed to assess both visual acuity (VA) and contrast sensitivity (CS). During testing, mice were placed on an elevated platform surrounded by four computer monitors displaying rotating sinusoidal gratings moving at a constant speed of 12°/s in either the clockwise or counterclockwise direction. For visual acuity assessment, the spatial frequency of the gratings was increased while contrast and speed were held constant. The highest spatial frequency that elicited a head-tracking response was recorded as the VA threshold, with higher values indicating better visual acuity. For contrast sensitivity assessment, gratings were presented at a fixed spatial frequency and speed while contrast was gradually reduced. Contrast sensitivity was defined as the lowest contrast level that still evoked a tracking response and was expressed as Michelson contrast: (L_max_ - L_min_)/(L_max_ + L_min_), where L_max_ and L_min_ represent the maximum and minimum luminance values of the grating, respectively. Measurements were obtained for both clockwise and counterclockwise stimulus rotations and subsequently averaged to generate a composite measure of visual performance.

**High-resolution diffusion MRI**

High-resolution in vivo diffusion MRI was performed using a 9.4 Tesla Bruker 94/30 system (Billerica, MA, USA) with a 30 cm bore and maximum gradient strength of 660 mT/m on each axis. Diffusion-weighted images (DWI) were acquired using a multi-shot 2D echo-planar imaging (EPI) pulse sequence with the following parameters: matrix size = 100 × 86 × 54, field of view (FOV) = 15 × 12.9 mm², in-plane resolution = 150 μm, slice thickness = 0.3 mm, echo time (TE) = 27 ms, repetition time (TR) = 3500 ms, and b-values of 600, 1200, 1800, and 3000 s/mm² with 4 non-diffusion-weighted (b0) volumes.

Prior to analysis, acquired DWI data were processed through a standardized preprocessing pipeline. Raw DWI data were first denoised using the Marchenko-Pastur Principal Component Analysis (MP-PCA) method(*2*), followed by removal of Gibbs ringing artifacts using a local subvoxel-shift method(*3, 4*). EPI-induced geometric distortions, eddy current-induced distortions, and subject motion were corrected by nonlinear registration to structural T2-weighted images using symmetric normalization (SyN)(*5*). Bias field correction was subsequently applied using the N4 algorithm implemented in ANTs. Brain masks were estimated and manually verified. The diffusion-weighted volumes were fitted to the tensor model to calculate diffusion tensor metrics, including Fractional Anisotropy (FA), Axial Diffusivity (AD), Radial Diffusivity (RD), and Mean Diffusivity (MD)(*6*). The data were fitted to kurtosis tensors to calculate diffusion kurtosis metrics, including Kurtosis Fractional Anisotropy (KFA), Axial Kurtosis (AK), Radial Kurtosis (RK), and Mean Kurtosis (MK)(*7*). Neurite Orientation Dispersion and Density Imaging (NODDI) modeling was carried out using the AMICO toolbox, which separates the contribution of neurite geometry from that of the surrounding extracellular space. This produced three indices for each voxel; neurite density index (NDI), orientation dispersion index (ODI), and free water fraction (FWF), the last of which accounts for isotropic fluid contamination such as CSF partial-volume effects(*8, 9*). For spatial normalization, the subject-space average B0 image was registered to our T2 brain atlas using ANTs, and the resulting transformation was used to warp atlas labels back into native subject space, enabling region-specific extraction of all diffusion metrics.

For tractography, A fully automated, unsupervised method was performed to obtain 5 tissue response functions (left OPT, left LGN, right OPT, right LGN, and right VC) using MRtrix3. Note that the left VC was excluded from analysis, as TBI caused damages in this region. A non-negativity constrained super-resolved spherical deconvolution method was used to compute the fiber orientation distribution (FOD)(*10*). The 5 tissue functions probabilistic tractography with 10 million streamlines and threshold 0.17 were first generated from those FODs(*11*), and the tracts were filtered using spherical-deconvolution informed filtering of tractograms (SIFT) to reduce reconstruction biases(*12*). Finally, the tracts of interest of tractography were kept between left OPT to left LGN, right OPT to right LGN, right LGN to right VC for observing the changes of fiber connection(*13*).

**Tissue processing**

The animals were deeply anesthetized and then perfused transcardially with cold saline, followed by a fixative containing 4% paraformaldehyde (PFA) in PBS. The brains and eyes were removed, post fixed overnight in 4% PFA, and then cryoprotected for 48 h in 30% sucrose. Serial 30 μm thick coronal or sagittal sections were cut using a cryostat (LeicaCM 1950, Teaneck, NJ), and stored at -20 °C. The sections were then processed for immunohistochemical analysis.

**Retinal flat mounting and RGC quantification**

Retinas from both ipsilateral and contralateral eyes were harvested at approximately 6 weeks (Day 44) post-injury. Immediately after collection, tissues were placed in ice-cold PBS and then fixed in 4% paraformaldehyde. Fixed retinas were incubated for five overnight periods with rabbit anti-RBPMS antibody, a retinal ganglion cell marker, at a 1:200 dilution (cat. no. AB194213, Abcam, Cambridge, UK). Following primary antibody incubation, tissues were labeled with Alexa Fluor 647-conjugated donkey anti-rabbit secondary antibody (cat. no. A31573, Invitrogen, Carlsbad, CA). DAPI was added during the final wash to counterstain nuclei. After immunostaining, eyecups were radially cut into a petal-like configuration and flat-mounted on slides using 0.45 μm black mixed cellulose ester membrane filters. Coverslips were applied with Aqua-Poly/Mount mounting medium. Confocal images were acquired using a Zeiss LSM700 microscope equipped with a 40×/1.3 NA oil-immersion objective. For each retina, eight z-stack images were collected from four retinal quadrants, including both central and peripheral regions. Maximum-intensity projections were generated from the z-stacks, and RBPMS-positive retinal ganglion cells were manually quantified.

**Cresyl Violet Staining Nissl**

Briefly, frozen coronal sections were mounted on Superfrost plus slides (Fisher Scientific, Pittsburgh, PA, USA) and stained using 0.1% cresyl violet solution for 20 minutes. Stained sections were rinsed in double-distilled water quickly and were immersed in 95% ethanol for 2 minutes. Then, the sections were dehydrated in 100% ethanol and were cleared in 100% xylene twice (5 minutes each time). Finally, the sections were mounted with DPX for imaging.

**Cortical Cavity Volume Measurement**

Briefly, whole sets of 1-in-6 series of 30-μm-thick brain coronal sections (180 μm apart) were stained with cresyl violet to show the spare cortex. Then, boundary contours of the contralateral and ipsilateral spare cortex were drawn using Photoshop (acrobat). The enclosed volume within the contours was measured, and the percent cortex of the cavity was calculated with the following formula: percentage of the cortical cavity = (contralateral cortex volume - ipsilateral spare cortex volume)/(contralateral cortex volume) × 100%(*14*).

**Immunofluorescence Staining.**

Immunofluorescence staining was carried out according to our previous reports(*15*). In brief, sections were rinsed in PBS 3 times (5 minutes each) and incubated in blocking buffer (0.1%Triton X-100, 1% bovine serum albumin, and 5% normal serum in PBS) for 1 hour at room temperature. Then, primary antibodies were added to blocking buffer and incubated with sections overnight at 4 °C. After 3 washes with PBS (5 minutes each), sections were incubated with the secondary antibody for 1 hour at room temperature and then treated with 4′,6-Diamidino-2-phenylindole (DAPI) for 2 minutes. Finally, sections were washed 3 times with PBS (5 minutes each), dried, and mounted using Fluoromount G (Southern Biotech, Birmingham,AL, USA). Primary antibodies and their final concentrations were as follows: NeuN antibody (1:1000, rabbit, Abcam, AB177487), synaptophysin antibody (1:1000, mouse, Millipore, MAB5258), GFAP antibody (1:1000, chicken, Abcam, AB4674), CD68 (1:500, rabbit, Abcam, AB213363). Secondary antibodies from Jackson ImmunoResearch Laboratories Inc. were all applied at the same dilution of 1:1000.

**Fluorescence intensity measurement and cell counting**

The intensity of synaptophysin or GFAP signal was measured using ImageJ software (version 1.47; NIH, Bethesda, MD). Briefly, images with synaptophysin-positive signal or GFAP signal were captured using a Zeiss inverted microscope (Zeiss Axio 200, Oberkochen, Germany). Three free sampling areas in injured cortex (epicenter) or contour of scar formation area (within 600 μm to the edge of cavity) was drawn using ImageJ software and the mean intensity of synaptophysin or GFAP signal within the contour was measured. The average of mean intensity from five mice was used for comparison. The NeuN-positive neurons in spared vision cortex were counted as described in our previous report(*16*). The contours of measuring areas (the block of injured cortex with 600 μm wide from edge of the cavity) were drawn. The NeuN-positive cells were quantified using an optical fractionator (MBF Bioscience, Williston, VT), and the data from five mice was averaged and presented. All CD68-positive cells were counted within injured cortex and the data from five mice was averaged and presented.

**Rotarod and elevated plus maze (EPM) tests**

Motor deficit and impulsiveness are common manifests after TBI, which can be commonly evaluated using the rotarod test or elevated plus maze (EPM) test, respectively. The rotarod test was conducted first to evaluate the effects of WAY treatment on balance and coordination of mice post-TBI versus sham as described in our previous report(*16*). The rotating rod was 3 cm in diameter and divided by flanges in five compartments to allow testing of up to five mice simultaneously. The animals had to walk on the rotating rod from the lowest speed to gradually increased speed (0-30 rpm within 90 s) and the time until the mouse fell from the rod (latency to fall) was recorded. A total of 4 trials were performed consecutively for each time point. Following the rotarod test, the EPM test was performed to assess anxiety-related behavior of mice as previously described(*17*). The EPM apparatus consists of a cross-shape platform with two oppositely positioned closed arms with high walls, two oppositely positioned open arms, and a small center area. The EPM apparatus was placed at 0.5 m from the ground. The EPM test is based on the natural aversion of rodents to open spaces. Mice were allowed to start at the maze center facing one of the open arms to explore the maze for a specific amount of time (10 min). As mice freely explored the maze, their behavior was recorded by a video camera mounted above the maze. Mouse behavioral metrics including total distance traveled and time spent in each arm were analyzed using ANY-maze software (Stoelting, Wood Dale, IL). The preference for being in open arms over closed arms (expressed as moving distance and time spent in the open arms) was calculated to measure anxiety-related behavior. Three groups with 7-10 mice for each were used for behavior tests including: sham, TBI, and TBI + WAY.

**Statistics and reproducibility**

Each animal was treated as an independent biological replicate. Sample sizes were based on group sizes commonly used in comparable CCI-TBI studies(*16, 18*). No animals or data points were excluded from analysis except if the animal is dead or the data contains statistical outliers during the longitudinal experiments. Animal group identity was masked by an independent researcher, and optokinetic, electrophysiological, and dMRI data acquisition and analysis were performed by separate experimenters blinded to treatment group.

The choice of statistical test was guided by the underlying experimental design rather than applied uniformly, as detailed below. For experiments in which three groups (Sham, TBI, and TBI+WAY) were compared across a single independent variable and the biological question concerned an overall group effect, one-way ANOVA with Tukey's post-hoc correction for multiple comparisons was used (Figure. 2B, C; Figure. 5E, I, M; Figure. 6K; Figure. 7B, D, E). This approach controls the family-wise error rate across all pairwise comparisons within a group of three or more. For experiments in which the biological question was restricted to two specific groups, rather than an comparison across all groups, unpaired two-tailed Student's t-tests were used for each comparison independently (Figure. 2F, G, I, J; Figure. 3B, C, E, F; Figure. 4D, E; Supplementary Figure. 1C, E; Supplementary Figure. 2C, E; Supplementary Table 1). This approach was applied specifically to hemisphere- and eye-specific electrophysiological and diffusion MRI measurements, where each eye or hemisphere constitutes an anatomically and functionally distinct dataset best evaluated independently rather than folded into a single compilation test. For the GFAP intensity analysis, in which immunoreactivity was quantified at multiple discrete distances from the lesion cavity edge, independent unpaired t-tests were performed at each distance bin (Figure. 6L), treating distance from the lesion as a defined spatial covariate rather than a level within a single ANOVA factor. For experiments involving two independent variables, namely treatment group and eye (ipsilateral vs. contralateral to injury), two-way ANOVA was used to resolve main effects of each variable as well as their interaction, followed by Tukey's post-hoc correction (Figure. 3I). All statistical analyses were performed in GraphPad Prism 11.1. Exact P values are reported in each figure panel. Data are presented as mean ± SEM (standard error of mean) unless otherwise indicated. Figures were assembled in Adobe Illustrator.

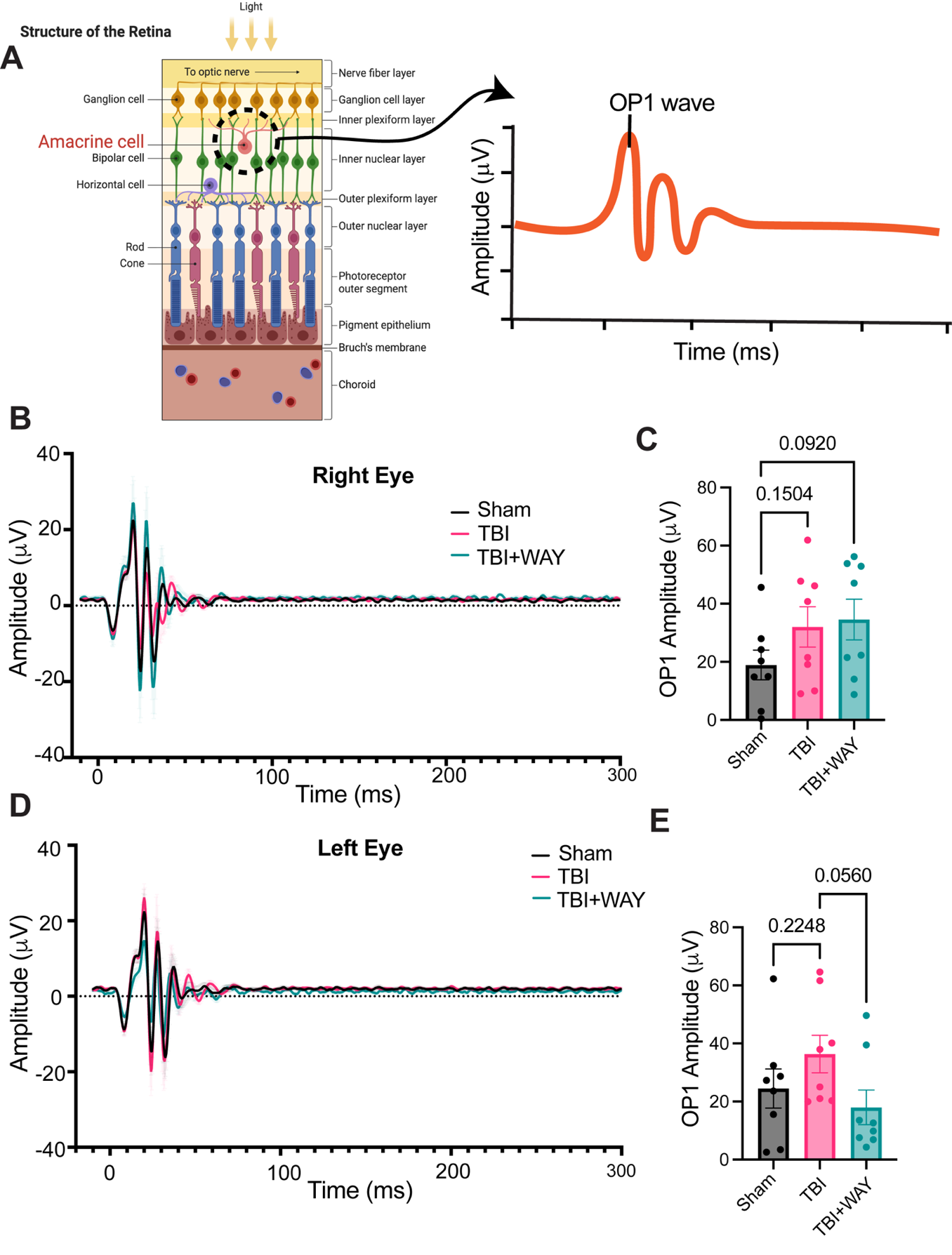
**Figures**

**Supplemental Figure 1. CCI-TBI induces heightened inner retinal output which is normalized by the WAY treatment.** (**A)** Schematic illustration the position of amacrine cells within the inner nuclear layer of the retina and a representative trace depicting the first oscillatory potential (OP1), reflecting retinal output driven by amacrine mediated inhibitory feedback onto RGCs. (**B, D)** Representative oscillatory potential (OP) traces from (B) right and (D) left eyes across experimental groups around 4 weeks (Day 25-26) after TBI; each trace represents the average per group. (**C, E)** Quantification of OP1 amplitude under the indicated conditions. Unpaired two-tailed *t*-test; *n* = 8 mice per group (Sham, TBI, TBI+WAY). Exact *P* values are indicated between compared groups in each panel.

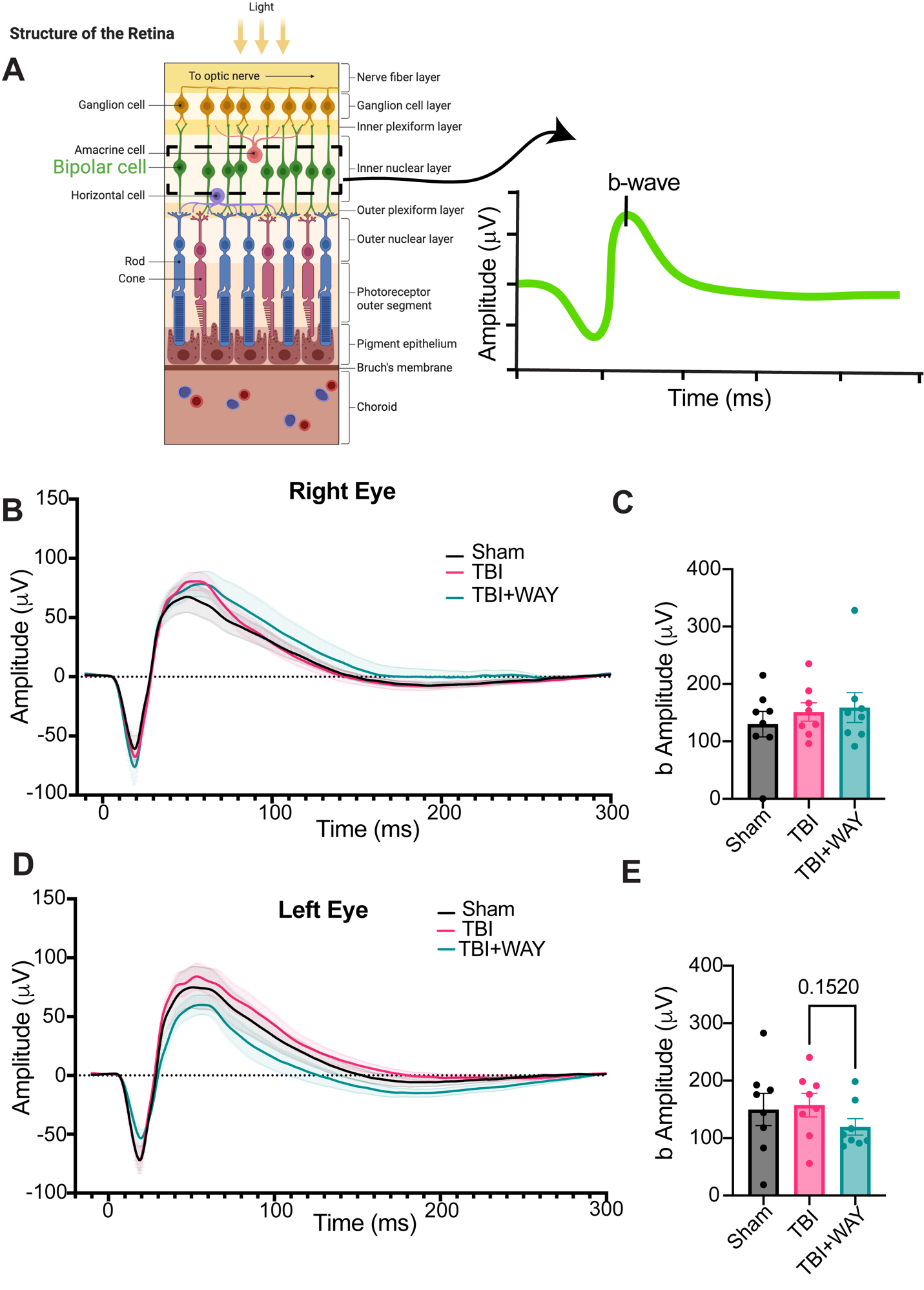

**Supplemental Figure 2. WAY treatment differentially modulates bipolar cell-driven retinal responses following CCI-TBI. (A)** Schematic illustration of the position of bipolar cells within the inner nuclear layer of the retina and a representative trace depicting the b-wave, reflecting depolarization of ON-bipolar cells in response to photoreceptor input and their excitatory drive onto downstream retinal circuits. **(B, D)** Representative b-wave ERG traces from (B) right and (D) left eyes approximately 4 weeks (Day 25-26) after TBI across experimental groups; each trace represents the average per group. **(C, E)** Quantification of b-wave amplitude under the indicated conditions. Unpaired two-tailed *t*-test; *n* = 8 mice per group (Sham, TBI, TBI+WAY). Exact *P* values are indicated between compared groups in each panel.

**Supplementary Table 1**

| **Metric** | **Sham (n=3)** | **TBI (n=5)** | **TBIWAY (n=6)** | **Sham vs TBI (p)** | **TBI vs TBI+WAY (p)** |
| --- | --- | --- | --- | --- | --- |
| **DTI** | | | | | |
| **Left_OPT_FA** | 0.428 | 0.489 | 0.47 | 0.024* | 0.020* |
| **Right_OPT_FA** | 0.362 | 0.457 | 0.445 | 0.030* | 0.774 |
| **Left_LGN_FA** | 0.353 | 0.365 | 0.373 | 0.52 | 0.601 |
| **Right_LGN_FA** | 0.372 | 0.409 | 0.388 | 0.033* | 0.091 |
| **Right_VC_FA** | 0.181 | 0.183 | 0.182 | 0.923 | 0.924 |
| **Left_OPT_MD** | 0.00037 | 0.00044 | 0.00043 | 0.088 | 0.624 |
| **Right_OPT_MD** | 0.00051 | 0.00048 | 0.00046 | 0.148 | 0.384 |
| **Left_LGN_MD** | 0.00048 | 0.0005 | 0.00049 | 0.31 | 0.581 |
| **Right_LGN_MD** | 0.00049 | 0.00047 | 0.00047 | 0.011* | 0.605 |
| **Right_VC_MD** | 0.00038 | 0.00043 | 0.00041 | 0.026* | 0.478 |
| **Left_OPT_AD** | 0.00061 | 0.00072 | 0.00068 | 0.059 | 0.424 |
| **Right_OPT_AD** | 0.00072 | 0.00075 | 0.00072 | 0.048* | 0.261 |
| **Left_LGN_AD** | 0.00066 | 0.00068 | 0.00068 | 0.427 | 0.77 |
| **Right_LGN_AD** | 0.00069 | 0.00069 | 0.00066 | 0.561 | 0.114 |
| **Right_VC_AD** | 0.00047 | 0.00052 | 0.00051 | 0.024* | 0.492 |
| **Left_OPT_RD** | 0.00026 | 0.0003 | 0.0003 | 0.147 | 0.84 |
| **Right_OPT_RD** | 0.0004 | 0.00034 | 0.00033 | 0.06 | 0.745 |
| **Left_LGN_RD** | 0.00039 | 0.0004 | 0.00039 | 0.349 | 0.472 |
| **Right_LGN_RD** | 0.00039 | 0.00036 | 0.00037 | 0.010* | 0.51 |
| **Right_VC_RD** | 0.00034 | 0.00038 | 0.00037 | 0.033* | 0.471 |
| **DKI** | | | | | |
| **Left_OPT_AK** | 1.085 | 1.273 | 1.216 | 0.039* | 0.328 |
| **Right_OPT_AK** | 1.046 | 1.13 | 1.124 | 0.009* | 0.903 |
| **Left_LGN_AK** | 1.255 | 1.218 | 1.24 | 0.406 | 0.503 |
| **Right_LGN_AK** | 1.136 | 1.144 | 1.159 | 0.756 | 0.503 |
| **Right_VC_AK** | 0.835 | 0.847 | 0.864 | 0.599 | 0.491 |
| **Left_OPT_MK** | 1.159 | 1.371 | 1.228 | 0.141 | 0.125 |
| **Right_OPT_MK** | 1.059 | 1.209 | 1.159 | 0.003* | 0.414 |
| **Left_LGN_MK** | 1.172 | 1.112 | 1.145 | 0.111 | 0.506 |
| **Right_LGN_MK** | 1.127 | 1.016 | 1.015 | 0.085 | 0.988 |
| **Right_VC_MK** | 0.656 | 0.69 | 0.711 | 0.202 | 0.319 |
| **Left_OPT_RK** | 1.162 | 1.471 | 1.219 | 0.096 | 0.037* |
| **Right_OPT_RK** | 1.036 | 1.239 | 1.154 | 0.003* | 0.21 |
| **Left_LGN_RK** | 1.066 | 0.951 | 1.021 | 0.022* | 0.393 |
| **Right_LGN_RK** | 1.035 | 0.782 | 0.807 | 0.057 | 0.804 |
| **Right_VC_RK** | 0.52 | 0.56 | 0.58 | 0.387 | 0.464 |
| **Left_OPT_KFA** | 0.482 | 0.544 | 0.536 | 0.013* | 0.506 |
| **Right_OPT_KFA** | 0.428 | 0.513 | 0.508 | 0.028* | 0.887 |
| **Left_LGN_KFA** | 0.599 | 0.609 | 0.591 | 0.645 | 0.446 |
| **Right_LGN_KFA** | 0.571 | 0.647 | 0.639 | 0.014* | 0.752 |
| **Right_VC_KFA** | 0.458 | 0.491 | 0.48 | 0.233 | 0.662 |
| **NODDI** | | | | | |
| **Left_OPT_NDI** | 0.789 | 0.898 | 0.878 | 0.023* | 0.441 |
| **Right_OPT_NDI** | 0.784 | 0.856 | 0.841 | 0.010* | 0.674 |
| **Left_LGN_NDI** | 0.85 | 0.827 | 0.838 | 0.149 | 0.49 |
| **Right_LGN_NDI** | 0.824 | 0.822 | 0.827 | 0.883 | 0.644 |
| **Right_VC_NDI** | 0.562 | 0.574 | 0.579 | 0.56 | 0.78 |
| **Left_OPT_ODI** | 0.32 | 0.35 | 0.358 | 0.31 | 0.678 |
| **Right_OPT_ODI** | 0.361 | 0.345 | 0.346 | 0.281 | 0.982 |
| **Left_LGN_ODI** | 0.507 | 0.505 | 0.489 | 0.961 | 0.541 |
| **Right_LGN_ODI** | 0.458 | 0.444 | 0.466 | 0.336 | 0.086 |
| **Right_VC_ODI** | 0.51 | 0.526 | 0.524 | 0.354 | 0.888 |
| **Left_OPT_FWF** | 0.03977 | 0.05011 | 0.05474 | 0.316 | 0.657 |
| **Right_OPT_FWF** | 0.11132 | 0.08001 | 0.07515 | 0.045* | 0.694 |
| **Left_LGN_FWF** | 0.02103 | 0.02625 | 0.02322 | 0.304 | 0.69 |
| **Right_LGN_FWF** | 0.01363 | 0.00878 | 0.00924 | 0.304 | 0.873 |
| **Right_VC_FWF** | 0.00211 | 0.00268 | 0.00269 | 0.658 | 0.995 |

**Supplementary Table 1: Diffusion MRI metrics across visual pathway regions following CCI-TBI and WAY treatment.** Mean values for DTI (FA, MD, AD, RD), DKI (AK, MK, RK, KFA), and NODDI (NDI, ODI, FWF) metrics in the left and right optic tract (OPT), left and right lateral geniculate nucleus (LGN), and right visual cortex (VC) across sham, TBI, and TBI+WAY groups, along with p-values between Sham vs TBI and TBI vs TBI+WAY. Unpaired two-tailed t-test; n (Sham) = 3, n (TBI) = 5, n (TBI+WAY) = 6 mice per group. *P < 0.05.

**Author’s contributions:**

A.D. conceived the study, supervised all aspects of the project, designed the experiments, interpreted the data, and wrote the manuscript. S.D. designed and performed experiments, analyzed and interpreted the data, and contributed to manuscript preparation. L.K.P. and X.G. performed the TBI procedures, elevated plus maze and rotarod studies, brain histology and imaging, analyzed the data, and contributed to manuscript preparation. N.W. and J.C. processed, analyzed, and interpreted the magnetic resonance imaging data and wrote the corresponding sections of the manuscript. All authors reviewed and approved the final manuscript.

**Acknowledgements:**

We thank Dr. Yu-Chien Wu and Erin E. Jarvis at the IU In Vivo Imaging Core for their assistance with MRI sample preparation and imaging. We thank the LARC Small Animal Tech Service for their help with ketamine/xylazine injections during fVEP recordings. We thank Michelle L. Surma for her assistance with animal identity blinding for PhNR recordings, optokinetic response testing and for her daily support with laboratory maintenance.

**REFERRENCES**

1.	J. Romine, X. Gao, J. Chen, Controlled cortical impact model for traumatic brain injury. *J Vis Exp*, e51781 (2014).
